# Host specialization in microparasites transmitted by generalist vectors: insights into the cellular and immunological mechanisms

**DOI:** 10.1101/2020.11.13.380550

**Authors:** Yi-Pin Lin, Danielle M. Tufts, Alan P. Dupuis, Matthew Combs, Ashley L. Marcinkiewicz, Andrew D. Hirsbrunner, Alexander J. Diaz, Jessica L. Stout, Anna M. Blom, Klemen Strle, April D. Davis, Laura D. Kramer, Maria A. Diuk-Wasser

**Author notes:** Contributed equally to this work (Yi-Pin Lin, Danielle M. Tufts). correspondence: Yi-Pin Lin, Ph.D., Division of Infectious Disease, Wadsworth Center, New York State Department of Health, 120 New Scotland Ave., Albany, NY 12047, correspondence: Maria A. Diuk-Wasser, Ph.D., Department of Ecology, Evolution, and Environmental Biology, Columbia University, 1200 Amsterdam Ave., New York, NY 10027.

## Abstract

Host specialization is an ecological and evolutionary process by which a pathogen becomes differentially adapted to a subset of hosts, restricting its host range. For parasites transmitted by generalist vectors, host specialization is not expected to evolve because of the decreased survival of those parasites in inadequate hosts. Thus, parasites may develop adaptation strategies, resulting in host specialization. The causative agents of Lyme disease are multiple species of bacteria, *Borrelia burgdorferi* sensu lato species complex (*Bb*sl), and are suitable for examining host specialization as birds and rodents were found to carry different species of these bacteria. Debate exists on whether host specialization occurs among these strains within a particular species of *Bb*sl, such as *B. burgdorferi* sensu stricto (*Bb*ss). Current evidence supports some *Bb*ss strains are widespread in white-footed mice but others are in non-rodent vertebrates, such as birds. To recapitulate specialization in the laboratory and define the mechanisms for host specialization, we introduced different genotypes of *Bb*ss via tick transmission to American robins and white-footed mice, the Lyme disease reservoirs in North America. Among these strains, we found distinct levels of spirochete presence in the bloodstream and tissues and maintenance by these animals in a host-dependent fashion. We showed that the late stage persistence of these strains largely corresponds to bacterial survival at early infection onsets. We also demonstrated that those early survival phenotypes correspond to spirochete adhesiveness, evasion of complement-mediated killing in sera, and/or not triggering high levels of pro-inflammatory cytokines and antibodies. Our findings thus link host competence to *Bb*ss with spirochete genotypic variation of adhesiveness and inducing/escaping host immune responses, illuminating the potential mechanisms that dictate host specialization. Such information will provide a foundation for further investigation into multi-disciplinary processes driving host specialization of microparasites.

**AUTHOR SUMMARY:** Host specialization arises when microparasites adapt to a subset of available hosts, restricting the host ranges they can infect. The mechanisms and selective pressures for the evolution of host specialization remain unclear. The causative agent of Lyme disease (LD), the bacteria species complex of *Borrelia burgdorferi* sensu lato, is adapted to different vertebrates. However, whether such a differential host adaption also applies to each genotype within the same species is under debate. Further, the mechanisms that drive such host specialization are unclear. We thus introduced three genotypes of one LD bacteria species *(B. burgdorferi* sensu stricto) individually via tick bite to American robins and white-footed mice, the most common LD reservoirs in North America. We found that these genotypes differed in the persistent maintenance by those reservoirs and occurred in a host-specific fashion. The ability of those bacteria for long-term maintenance was linked with their capability to attach to cells and a lack of induction of high levels of immune responses at early infection onsets. This work demonstrates the potential mechanisms that dictate host specialization of LD bacteria circulating in natural populations. Such information will pave the road to define the molecular, ecological, and evolutionary determinants that drive host-microparasite interactions.

## INTRODUCTION

The range of hosts a parasite can infect is arguably one of the most important properties of a parasite because it can determine, among other things, whether a parasite can survive the extinction of a host species and whether it can become established and spread following its introduction to a new area (1, 2). Host specialization is defined as the ecological and evolutionary process by which a pathogen becomes differentially adapted and thus restricts its host range to a subset of potential hosts. For vector-borne microparasites, such a process is expected to evolve when vectors are specialized to competent hosts, leading to microparasite amplification. However, it is much less obvious how and when host specialization in a vector-borne microparasite can occur and when vectors are host-indiscriminate. Given the significant cost incurred when microparasites are inoculated into incompetent hosts, the evolution of host specialization is only expected to occur when it provides a significant selective advantage for the microparasite to overcome cellular or immunological barriers to infection.

The spirochetes of the *B. burgdorferi* sensu lato (s.l.) species complex, agents of Lyme disease, represent an ideal system to investigate the tradeoffs involved in the evolution of host specialization. *B. burgdorferi* s.l. is maintained in an enzootic cycle between generalist ticks of the *Ixodes ricinus* complex and reservoir hosts, including small and medium-sized mammals, birds, and reptiles (3–5). Despite being transmitted by the same generalist vector, some *B. burgdorferi* s.l. genospecies in Europe are almost exclusively associated with a host taxa (e.g. *B. afzelii* in rodents; *B. garinii* in avian hosts). In contrast, *B. burgdorferi* sensu stricto (hereafter *B. burgdorferi*) in North America is considered a host generalist as it has been isolated from multiple classes of vertebrate animals including mammalian and avian hosts (3, 6–8). However, laboratory infection studies indicate that the fitness of *B. burgdorferi* strains varies in different hosts (9, 10).

In natural populations, weak associations were also found between hosts and particular *B. burgdorferi* genotypes, defined by polymorphic markers, such as *ospC* or 16S-23S rRNA intergenic spacer type (RST) (11–13). An intriguing possibility is that the partially and regionally constrained host associations observed in *B. burgdorferi* genotypes represent an incipient evolutionary process of host specialization (4, 12, 14–17). That is, *B. burgdorferi* may be on an evolutionary path to diversify into host specialized genospecies, similar to those within the *B. burgdorferi* s.l. species complex in Europe. Nonetheless, the molecular processes underlying such diversification and the extent to which they might drive genome-wide diversification in *B. burgdorferi* remain under debate.

Multiple-niche polymorphism (MNP), or diversifying selection, has been proposed as a balancing selection mechanism maintaining the diversity of *B. burgdorferi* genotypes through adaptation to host ‘niches’ (11, 12, 18, 19). This model was applied to the maintenance of the polymorphism in the outer surface protein C (OspC), one of the most diverse Lyme borreliae antigens that is heavily targeted by the vertebrate immune system (20–22). Alternatively (or concurrently), the OspC polymorphism could be maintained in a host-independent manner, through negative frequency-dependent selection. In this form of balancing selection, rare *B. burgdorferi OspC* genotypes to which few hosts have been exposed would have a selective advantage over the more common genotypes to which many hosts have mounted an adaptive immune response; thus maintaining diversity in the population (14, 17, 23–25). Theoretical studies have examined patterns of OspC diversity predicted by the different proposed eco-evolutionary mechanisms in different ecological settings (25–28), but a mechanistic understanding of the cellular and immunological mechanisms mediating strain-host interactions is critical to disentangle the multiple co-occurring selective pressures.

In this study, we simultaneously assessed cellular and immunological mechanisms that mediate strain-host interactions using three *B. burgdorferi* strains with variable *ospC* genotypes, as well as the North American rodent and avian reservoir hosts, American robins (hereafter robins) and white-footed mice (*Peromyscus leucopus*), respectively. We aimed to determine which cellular and immunological mechanisms mediate fitness variation in strain-host species pairs. Specifically, we identified genotypic differences across *B. burgdorferi* interacting with hosts in (1) the role of complement inhibition and inflammation induction; (2) the role of bacterial adhesion; (3) the role of antibodies in driving late stage persistence in specific hosts and (4) transmission efficiency to larval ticks. By characterizing these mechanisms, we aimed to identify plausible selective pressures that shape the composition of *B. burgdorferi* strain community, its diversity, and population dynamics.

## RESULTS

### *Borrelia burgdorferi* B31-5A4, 297, and cN40 differed in their adhesiveness to fibroblasts, *ex vivo* cytokine induction, and infection establishment in robins and white-footed mice

To compare the capability of genotypically distinct *B. burgdorferi* to initiate infection in reservoir hosts, the cloned *B. burgdorferi* strains B31-5A4, 297, and cN40 belonging to different genotypes (Table S1) and displaying similar growth rates *in vitro* (Fig. S1) were used in this study. Bacterial burdens at inoculation sites of skin and in blood were determined at 1 day after each of these strains was intradermally introduced into robins and white-footed mice. We did not detect any of these strains in robin blood (Fig. S2A), in agreement with no hematogenous dissemination at such an early time point (29). However, we found that B31-5A4 and cN40 showed significantly higher spirochete burdens at the initial injection sites, compared to mock-infected robins (Fig. 1). The injection sites from only two out of four 297-inoculated robins had spirochete burdens higher than detection limits (10 bacteria per 100ng of DNA from tissues), leading to non-significant differences in the levels of colonization from mock-infected robins (Fig. 1A). In white-footed mice infections, we did not observe spirochete burdens higher than detection limits of any of these strains in the blood of white-footed mice (Fig. S2B). Nonetheless, the burdens of B31-5A4 and 297 were significantly higher than those from mock-infected individuals at the inoculation site (~10^2^ bacteria per 100ng of DNA). Only two of five white-footed mice had cN40 bacterial loads greater than the detectable threshold limit, resulting in non-significant differences in burdens from mock-infected mice (Fig. 1B). These findings indicate B31-5A4, 297, and cN40 have the capacity to establish infection selectively in robins and white-footed mice.

**Figure 1.**
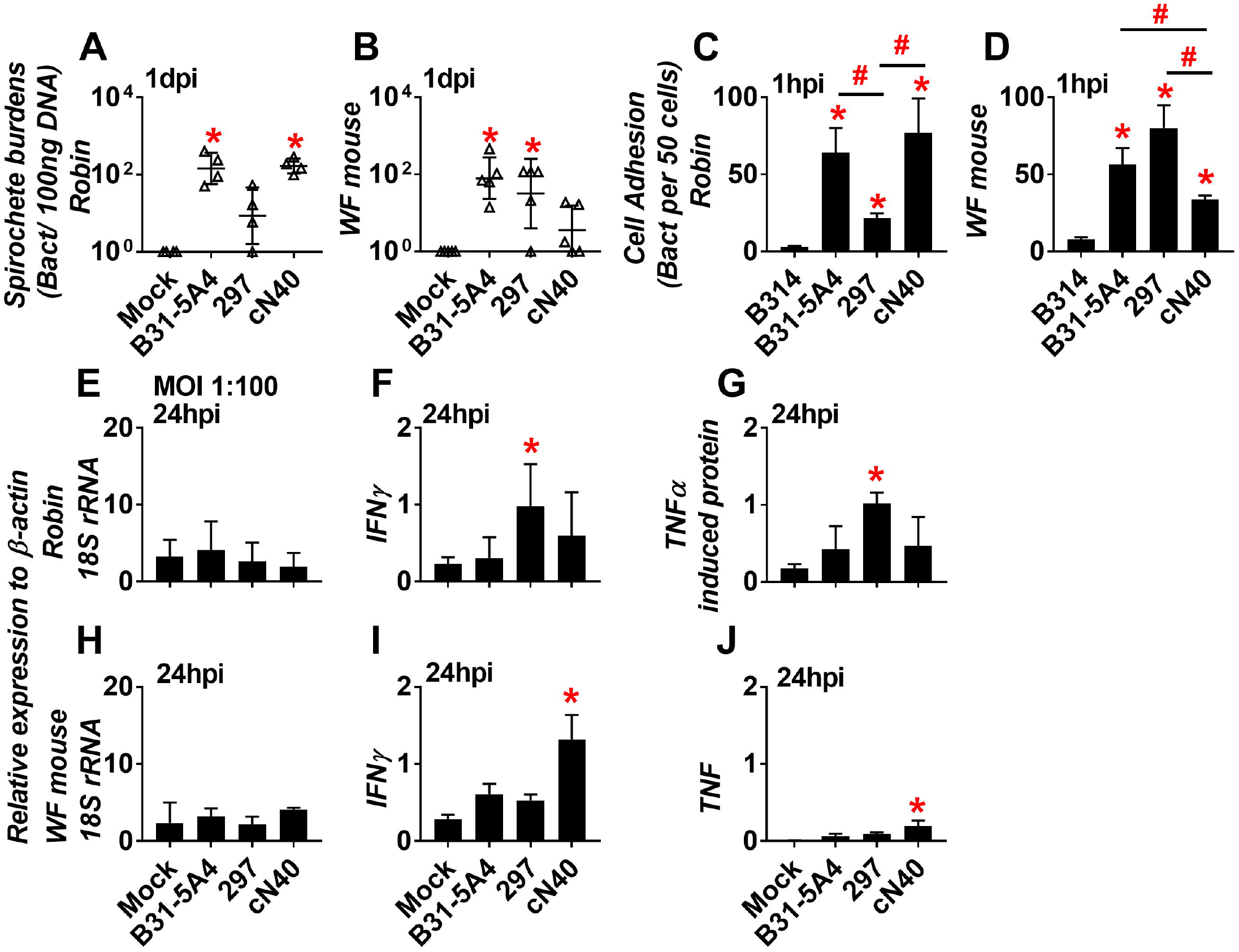
*Borrelia burgdorferi* B31-5A4, 297, and cN40 differed in early colonization in robins and white-footed mice and adhesion and cytokine triggering in these animals-derived fibroblasts. **(A and B): (A)** American robins and **(B)** white-footed (WF) mice were intradermally inoculated with 10^4^ *B. burgdorferi* B31-5A4, 297, or cN40, or with BSK-II medium without rabbit sera as mock infection (“Mock”). The inoculation site of skin from these robins and mice were collected at 1-day post infection (1dpi) to determine bacterial burdens by qPCR. The bacterial loads in the tissues or blood from robins or mice were normalized to 100 ng total DNA. Shown are the geometric mean ± geometric standard deviation of bacterial burdens in those tissues from four robins or mice per group. Significant differences (P < 0.05) in the spirochete burdens (normalized qPCR quantity values) relative to the mock infected group are indicated (“*”). **(C and D):** *B. burgdorferi* B31-5A4, 297, cN40, or B314 (negative control) (2 ×10^6^ spirochetes) were incubated with fibroblasts (2 ×10^5^ cells) from **(C)** American robins and **(D)** white-footed (WF) mice for 1 h. We mixed those cells with FITC-conjugated goat anti-*B. burgdorferi* polyclonal antibodies and visualized the spirochetes after fixation. Additionally, DAPI was incubated with these cells to localize the nuclei from fibroblasts. The levels of spirochete attachment were evaluated by counting the number of bacteria per 50 cells under fluorescence microscopy described in the section “Materials and Methods.” Each bar represents the mean of four independent determinations ± standard deviation. Asterisks indicate significant differences (P < 0.05) in spirochetal attachment relative using normalized qPCR quantity values to strain B314. **(E to J):** Fibroblasts (2 ×10^5^ cells) from **(E to G)** American robins and **(H to J)** white-footed (WF) mice were incubated for 24h with *B. burgdorferi* B31-5A4, 297, or cN40 (2 ×10^7^ of spirochetes for spirochete to cell ration (MOI) at 1:100). Cell media-treated fibroblasts were included as control (“Mock”). After the RNA was extracted from these cells, the expression levels of the genes encoding IFN-γ, TNF, or TNFα-induced protein and the constitutively expressed gene, actin, and 18S rRNA, from robins and white-footed mice were determined using quantitative reverse transcription polymerase chain reaction. The expression levels of the genes encoding **(E and H)** 18S rRNA, **(F and I)** IFNγ, **(G)** TNFα-induced protein, and **(J)** TNF are presented by normalizing to the expression levels of the gene encoding actin. Each bar represents the mean of three independent determinations ± SD. The asterisk (“*”) indicates significant differences *(P* < 0.05;) in the normalized expression levels of the gene encoding IFN-γ, TNF, and TNFα-induced protein in fibroblasts treated with indicated spirochete strains relative to those in mock-treated cells.

As the burdens of *B. burgdorferi* during infection initiation can be attributed to differences in spirochete attachment to cells or clearance by host inflammatory responses, we incubated B31-5A4, 297, and cN40 with fibroblasts isolated from robin or white-footed mouse for 1 h and determined the levels of bacterial attachment using fluorescent microscopy (Fig. S3). A nonadhesive *B. burgdorferi* strain B314 was also included as control (Table S1). Compared to B314, we found that B31-5A4, 297, and cN40 more efficiently bind to the fibroblasts from robins (Fig. 1C) or white-footed mice (Fig. 1D). Whereas B31-5A4 and cN40 bound to robin fibroblasts at significantly higher levels compared to 297 (Fig. 1C), B31-5A4 and 297 attached to white-footed mouse fibroblasts at significantly greater levels compared to cN40 (Fig. 1D). To determine whether B31-5A4, 297, and cN40 trigger differential pro-inflammatory responses in fibroblasts from different reservoir origins, we measured the expression levels of the genes encoding pro-inflammatory cytokines, IFNγ and TNF, or the proteins related to these cytokines after incubating cells with each of the spirochete strains for 24h. The expression levels of housekeeping genes (18S rRNA and β-actin) were also included as controls. We found that the levels of expression for each of the genes in any spirochete-treated cells were not significantly different from the mock-treated cells at 1 to 10 of the spirochetes to cell ratio (Fig. S4). At 1 to 100 of spirochete to cell ratio, the B31-5A4- or cN40-treated robin fibroblasts expressed similar levels of the tested cytokines to the cells under mock treatment while 297-incubated cells had significantly greater expression of IFNγ and TNF-induced protein than mock-treated cells, respectively (Fig. 1F and G). Conversely, while white-footed mouse fibroblasts incubated with B31-5A4 or 297 expressed those cytokines indistinguishably from mock-treated cells, cN40-treated cells showed significantly higher expression of IFNγ and TNF (4.6- and 30.1-fold, respectively) than the cells under mock treatment (Fig. 1I and J). These results suggest that *B. burgdorferi* B31-5A4, 297, and cN40 differ in their fibroblast adhesiveness and cytokine induction in a host-dependent fashion.

### Ticks acquired different levels of *B. burgdorferi* B31-5A4, 297, and cN40 from robins and white-footed mice infected with each of these spirochete strains

To compare reservoir host competence to B31-5A4, 297, and cN40, we assessed spirochete transmission from host to feeding larvae. Unfed *I. scapularis* nymphs were allowed to feed on robins or white-footed mice until repletion. We subsequently measured bacterial loads in flat and fed nymphs and found indistinguishable burdens among these ticks (~10^4^ bacteria per nymph), indicating no differences of the ability for any tested strain to survive in flat or fed nymphs (Fig. S5).

We then placed *I. scapularis* larvae on robins or white-footed mice at different time points (Fig. 2A) to determine spirochete burdens in each larva and the percentage of spirochete-positive larvae per individual host (defined as percent positivity). The spirochete burdens in uninfected (control) robin-derived larvae were below the detection limit (10 bacteria per tick), resulting in zero spirochete positivity at 14, 28, 35, and 56 days post nymph feeding (dpf) (Fig. 2B to F and Table 1). At least 18% of larvae feeding on B31-5A4-, 297-, or cN40-infected robins were spirochete positive at all time points with greater than detection limits of bacterial loads, indicating the ability of robins to persistently maintain and transmit these strains (Fig. 2C to F and Table 1). The percent positivity of fed larvae carrying cN40 was significantly higher than those harboring 297 at all four time points and significantly greater than those carrying B31-5A4 at 28, 35, and 56 dpf (Fig. 2B and Table 1). Moreover, the cN40-infected larvae had significantly higher bacterial loads than the 297-infected larvae at all time points and significantly greater than those in B31-5A4-infected larvae at 28, 35, and 56dpf (Fig. 2C to F). These data suggest that the cN40 is maintained and transmitted more efficiently in robins than B31-5A4 and 297.

**Figure 2.**
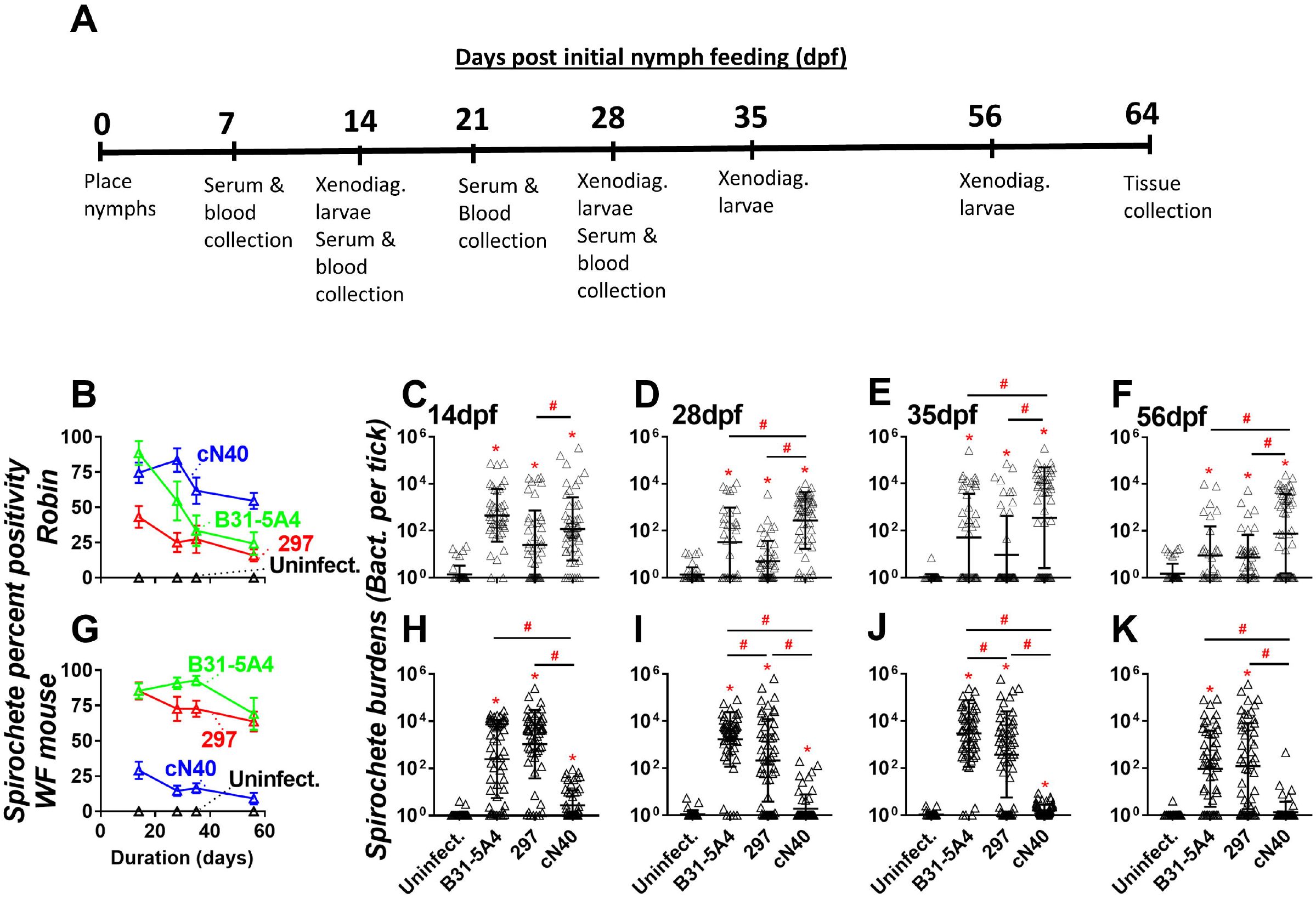
*Borrelia burgdorferi* B31-5A4, 297, and cN40 displayed strain-to-strain variation of xenodiagnostics acquisition from robins and white-footed mice. **(A)** The experimental timeline of infection. **(B to K):** *Ixodes scapularis* nymphs carrying *B. burgdorferi* B31-5A4, 297, or cN40, or naïve nymphs (Uninfect.) were allowed to feed to repletion on **(B to F)** 3, 4, 5, or 4 American robins, respectively, or **(G to K)** 5 white-footed (WF) mice per group. Approximately 100 larval ticks were placed on each robin at the time points indicated in Fig. 2A to feed till repletion. qPCR was used to determine spirochete burdens derived from 55 larvae feeding on *B. burgdorferi-* infected white-footed mice or 33, 44, 55, and 44 larvae feeding on robins fed on by nymphs carrying B31-5A4, 297, or cN40, or uninfected nymphs, respectively. **(B and F)** The larvae were considered xenodiagnostic positive if their spirochete burdens were greater than the threshold, the mean plus three-fold standard deviation of spirochete burdens in the uninfected group. Shown are the means ± SEM of percent positive larvae. **(C to F and H to K)** Shown are the geometric means ± geometric standard deviation of bacterial burdens in larvae that are allowed to feed on robins or white-footed (WF) mice at **(C and H)** 14, **(D and I)** 28, **(E and J)** 35, and **(F and K)** 56 days post nymph feeding (dpf). Significant differences (p < 0.05) in the spirochete burdens relative to larvae feeding on naïve robins or white-footed mice (“*”) or between different groups (“#”) are indicated.

**Table 1.** The number of positive xenodiagnostic larval ticks collected from robins and whitefooted mice.

| dpf<sup>a</sup> | Positive xenodiagnostic larval ticks acquired from |  |  |  |
| --- | --- | --- | --- | --- |
|  | Mice fed on by naïve nymphs | Mice fed on by nymphs carrying *B. burgdorferi* |  |  |
|  |  | B31-5A4 | 297 | cN40 |
| **American robins** |  |  |  |  |
| **14** | 0/44 | 39/44<sup>b</sup> | 19/44<sup>b,c</sup> | 41/55<sup>b</sup> |
| **28** | 0/44 | 18/33<sup>b,c</sup> | 11/44<sup>b,c</sup> | 46/55<sup>b</sup> |
| **35** | 0/44 | 11/33<sup>b,c</sup> | 12/44<sup>b,c</sup> | 34/55<sup>b</sup> |
| **56** | 0/44 | 8/33<sup>b,c</sup> | 7/44<sup>b,c</sup> | 30/55<sup>b</sup> |
| **White-footed mice** |  |  |  |  |
| **14** | 0/55 | 47/55<sup>b,c</sup> | 47/55<sup>b,c</sup> | 16/55<sup>b</sup> |
| **28** | 0/33 | 50/55<sup>b,c</sup> | 40/55<sup>b,c</sup> | 8/55<sup>b</sup> |
| **35** | 0/33 | 51/55<sup>b,c</sup> | 40/55<sup>b,c</sup> | 9/55<sup>b</sup> |
| **56** | 0/33 | 38/55<sup>b,c</sup> | 35/55<sup>b,c</sup> | 5/55<sup>b</sup> |

<sup>a</sup>Days post nymph feeding.
<sup>b</sup>Difference compared with larvae from the mice fed on by naïve nymphs by two-tailed Fisher test.
<sup>c</sup>Difference compared with larvae from the mice fed on by nymphs carrying strain cN40 by two-tailed Fisher test.

Similarly, we found bacterial burdens lower than detection limits in the larvae feeding on uninfected (control) mice, resulting in zero spirochete positivity (Fig. 2G to K, Table 1). At least 9% of the larvae feeding on B31-5A4-, 297-, or cN40-infected mice were positive throughout the experiment (Fig. 2G and Table 1). The bacterial loads from those mice were also statistically different from those derived from uninfected mice (Fig. 2H to K), except that larvae infected with cN40 had similar spirochete burdens to uninfected mouse-derived larvae at 56dpi. These data indicate the ability of white-footed mice to maintain and transmit each of these strains. We found significantly lower spirochete positivity in larvae infected with 297, compared to that in larvae infected with B31-5A4 at 28 and 35 dpf, respectively (p < 0.05, Fig. 2G, I, J and Table 1). Finally, larvae infected with cN40 showed more than significantly lower spirochete positivity (Fig. 2G and Table 1) and had significantly less bacterial burdens than B31-5A4- and 297-infected ticks throughout the experiments (Fig. 2H to K). These data suggest less capability of white-footed mice to maintain and transmit cN40 compared to B31-5A4 or 297.

### *Borrelia burgdorferi* B31-5A4, 297, and cN40 varied in their capability to trigger bacteremia and persistently colonize robins and white-footed mice

We next determined the ability of B31-5A4, 297, and cN40 to survive in the bloodstream of robins and white-footed mice. We found that robins infected with cN40 develop significantly greater levels of bacteremia at 7, 14, and 21dpf than the uninfected robins, which had spirochete burdens lower than the detection limits (10 bacteria per 100ng DNA, Fig. 3A to C). However, the burdens of that strain in the blood was no statistically different from those in uninfected robins at 28dpf (Fig. 3D). Robins infected with 297 had spirochete burdens in the blood no statistically different from uninfected robins at 7, 21 and 28 dpf but developed statistically greater bacterial loads at 14dpf (Fig. 3A to D). Though B31-5A4 was capable of inducing bacteremia in robins at 7 and 14dpf at the levels statistically greater than those in uninfected robins, the burdens of this strain in the blood were not statistically different from those in uninfected robins at 21 and 28dpf (Fig. 3A to D). These results indicate the ability of cN40 to induce long-lasting bacteremia in robins, compared to other strains, and 297 displayed delayed onsets of bacteremia. Similar to uninfected robins, uninfected white-footed mice did not exhibit bacteremia above the levels of detection throughout the experiment (Fig. 3E to H). At 21 and 28dpf, the spirochetes burdens in the blood from the mice infected with any of tested strains were no statistically different from those in uninfected mice (Fig. 3E and H). At 7 and 14 dpf, B31-5A4 and 297 triggered significantly higher levels of bacteremia than uninfected mice (Fig. 3E and F). However, the majority of mice infected with cN40 did not show detectable bacteremia (3/5 and 5/5 mouse blood had burdens below detection limits at 7 and 14dpf, respectively), resulting in no significant different bacterial loads from uninfected mice (Fig. 3E and F). These data suggest that cN40 is less capable of surviving in the white-footed mouse bloodstream compared to B31-5A4 and 297, in contrast to the results derived from robins.

**Figure 3.**
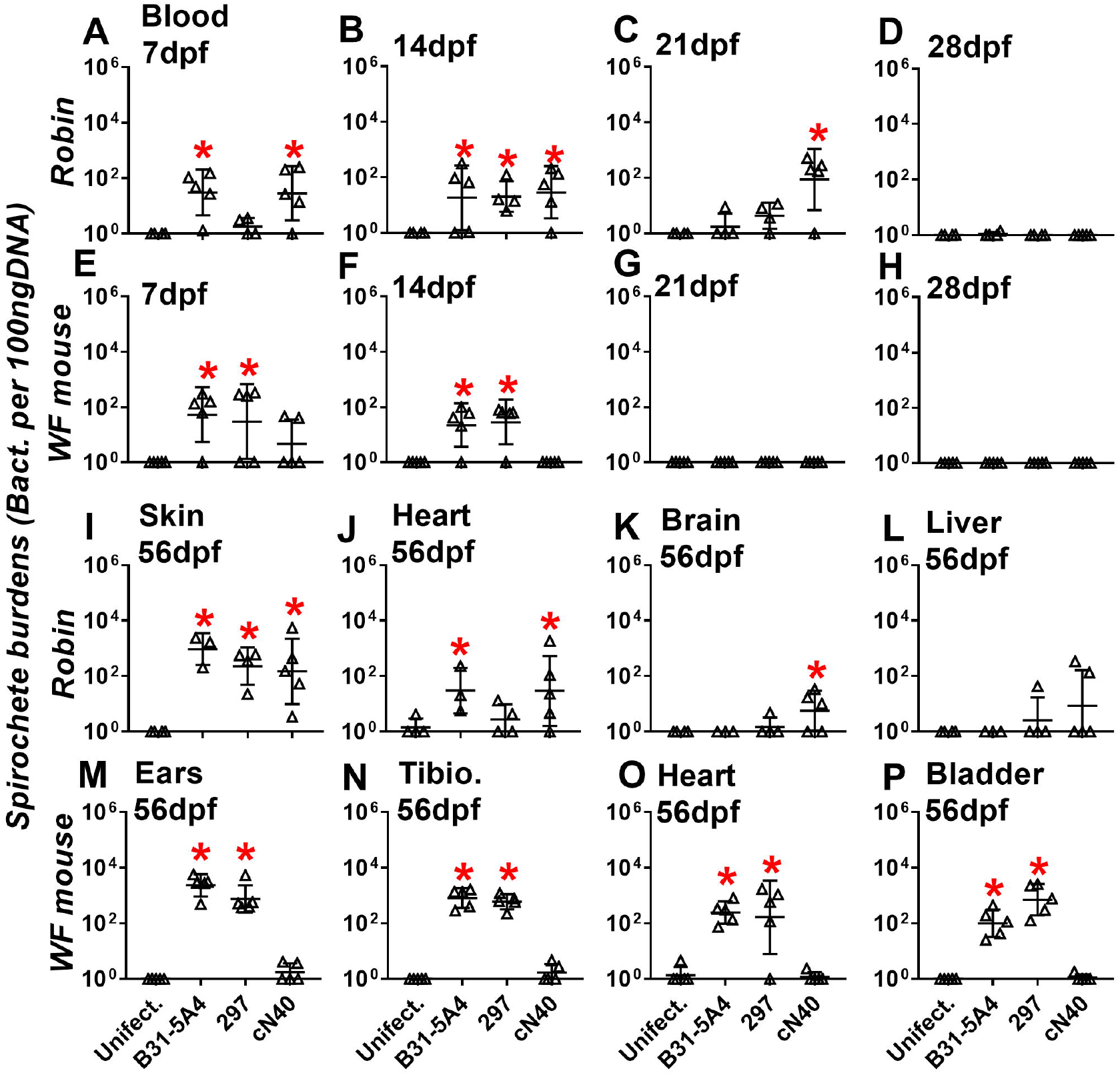
*Borrelia burgdorferi* B31-5A4, 297, and cN40 differed in their ability to induce bacteremia and colonize tissues in robins and white-footed mice. *Ixodes scapularis* nymphs carrying *B. burgdorferi* B31-5A4, 297, or cN40, or naïve nymphs (Uninfect.) were allowed to feed to repletion on **(A to D and I to L)** 3, 4, 5, and 4 American robins, respectively, or **(E to H and M to P)** 5 white-footed (WF) mice per group. The bacterial loads in the blood at **(A and E)** 7, **(B and F)** 14, **(C and G)** 21, and **(D and H)** 28 days post nymph feeding (dpf) and in **(I)** skin, **(J)** heart, **(K)** brain, and **(L)** liver of robins and **(M)** ears, **(N)** tibiofemoral joints (Tibio.), **(O)** heart, **(P)** bladder of white-footed mice at 64 days post nymph feeding (dpf) were determined by qPCR. The bacterial loads in blood were normalized to 100 ng total DNA. Shown are the geometric mean ± geometric standard deviation of indicated number of robins or white-footed mice. Significant differences (p < 0.05) in the spirochete burdens relative to robins or white-footed mice fed on by naïve nymphs (“*”) are indicated.

We also evaluated the ability of B31-5A4, 297, and cN40 to colonize robin and white-footed mouse tissues at 64dpf (Fig. 2A). Robins infected with each of these strains had significantly greater spirochete burdens in skin than uninfected robins, which exhibited bacterial loads below the detection limits (Fig. 3I). In addition, spirochetes were detected at significantly greater burdens in the heart and brain of cN40-infected robins and the heart of robins infected with B31-5A4, compared to respective tissues from uninfected robins (Fig. 3J and K). Majority of the 297-or cN40-infected robins (3 out of 4 and 3 out of 5 in 297- and cN40-infected robins, respectively) had burdens below the detection limits in livers, yielding no significantly different burdens from those in uninfected robins (Fig. 3L). Similarly, the burdens of B31-5A4 in the livers from spirochete-infected robins were no significantly different from those from uninfected robins (Fig. 3L). These data showed persistent skin colonization of strains B31-5A4, 297, and cN40 in robin, but the ability of each strain to colonize other tissues varied (Fig. 3I to L). We also measured the bacterial burdens at 64dpf in the tissues from white-footed mice infected with each of these strains. We detected spirochetes in ears, tibiofemoral joints, heart, and bladder from mice infected with B31-5A4 or 297 (~10^2^ to 10^3^ bacteria per 100ng of DNA from tissues). The bacterial burdens in tibiofemoral joints, heart, and bladder from B31-5A4- or 297-but not cN40-infected mice were significantly greater than those in uninfected mice (Fig. 3M to P). These findings demonstrated the ability of B31-5A4 and 297 but not cN40 to persistently colonize white-footed mice.

### Robin but not white-footed mouse complement differentiated the ability of *B. burgdorferi* B31-5A4, 297, and cN40 to survive in sera

To define the capability of B31-5A4, 297, and cN40 to evade complement-mediated killing, we evaluated the percent survival after incubation of these strains individually with the sera from uninfected robins or white-footed mice by counting the number of motile bacteria under microscopes. More than 95% of all tested strains including a serum sensitive control strain, *B. burgdorferi* B313 (30), survived in heat inactivated robin sera, in which the heat sensitive components such as complement were not functional (Fig. 4A and Table S1). Less than 16% of B313 remained motile, verifying the bactericidal activity of these robin sera (Fig. 4A). More than 85% of B31-5A4 and cN40 were alive, but only 50% of 297 was motile after incubated with robin sera (Fig. 4A). However, all strains survived at similar levels (greater than 80% of live spirochetes) in the sera pre-treated with OmCI, which inactivates robin complement by abolishing the lytic activity towards gram negative bacteria (Fig. 4B) (31). These results indicate that the 297 is more vulnerable to robin complement-mediated killing than B31-5A4 and cN40. We also determined the ability of spirochete strains to evade killing by white-footed mouse complement in the similar fashion. Greater than 95% of B313, B31-5A4, 297, and cN40 survived in heat inactivated white-footed mouse sera (Fig. 4C). Whereas only 29% of B313 remained motile after treated with these sera, more than 97% of other tested strains were alive in the untreated white-footed mouse sera (Fig. 4C). More than 93% of tested strains remained motile in white-footed mouse sera treated with CVF, which inactivates mammal complement (Fig. 4D) (32). These results indicate that all tested strains evade the killing by white-footed mouse complement at indistinguishable levels.

**Figure 4.**
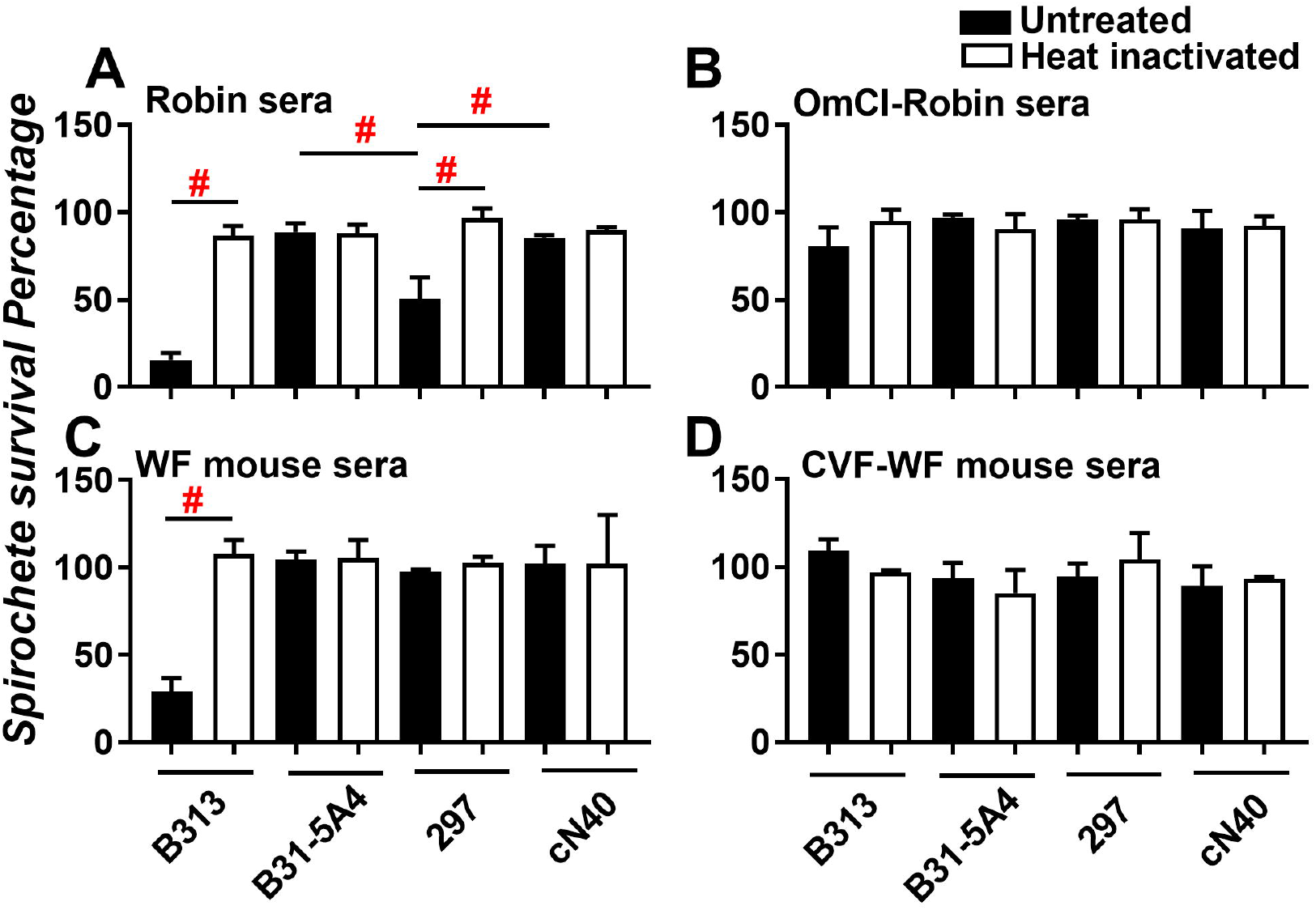
*Borrelia burgdorferi* B31-5A4, 297, and cN40 varied in their ability to survive in robin but not white-footed mouse sera. A high passage, non-infectious, serum sensitive *B. burgdorferi* strain B313 (“B313”) or *B. burgdorferi* B31-5A4, 297, or cN40 were incubated for 4 h with the sera from American robins at a final concentration of 40% in the **(A)** absence or **(B)** presence of 2 μM of OmCI. Each of these spirochete strains was also incubated for 4 h with the sera from white-footed (WF) mice at a final concentration of 40% in the **(C)** absence or **(D)** presence of 2 μM of CVF. The above-mentioned sera were also heat-inactivated and included as controls. The number of motile spirochetes was assessed microscopically. The percentage of survival for *B. burgdorferi* was calculated using the number of mobile spirochetes at 4 h post incubation normalized to that prior to the incubation with serum. The assays were performed at three independent occasions; within each experiment, samples were run in triplicate, and the survival percentage for each experiment was calculated by averaging the results from triplicate runs. The result shown here are the average ± standard deviation of the survival percentage from three independent experiments. Significant differences (P < 0.05) of the percent survival of spirochetes between groups are indicated (#).

### Robins and white-footed mice generated different levels of pro-inflammatory cytokines in response to early infection of *B. burgdorferi* B31-5A4, 297, and cN40

We compared the levels of pro-inflammatory cytokines, IFNγ and TNFα, in the sera derived from robin- and white-footed mice at different times during the infection. We found no significant different levels of IFNγ and TNFα in robins prior to the infection among different infection groups (Fig. 5A and E). At 7dpf, B31-5A4- or cN40-infected robins were not significantly different in those cytokines compared to uninfected robins, 297-infected robins produced significantly greater levels of IFNγ and TNFα than those uninfected birds, respectively (Fig. 5B and F). At 14dpf, the levels of these cytokines from those robins are no different from those in uninfected robins. In contrast to robins at 7dpf, cN40-infected mice displayed significantly higher levels of these cytokines than uninfected mice at this time point while 297-infected mice had no significantly different in the levels of those cytokines, compared to uninfected mice (Fig. 5J and N). Note that B31-5A4-infected mice had significantly greater levels of IFNγ (Fig. 5J) but were not significantly different from levels of TNFα at 7dpf (Fig. 5N), compared to uninfected mice. These data indicate differences of pro-inflammatory cytokine induction by each of these spirochete strains, particularly at early stages of robin and white-footed mouse infection.

**Figure 5.**
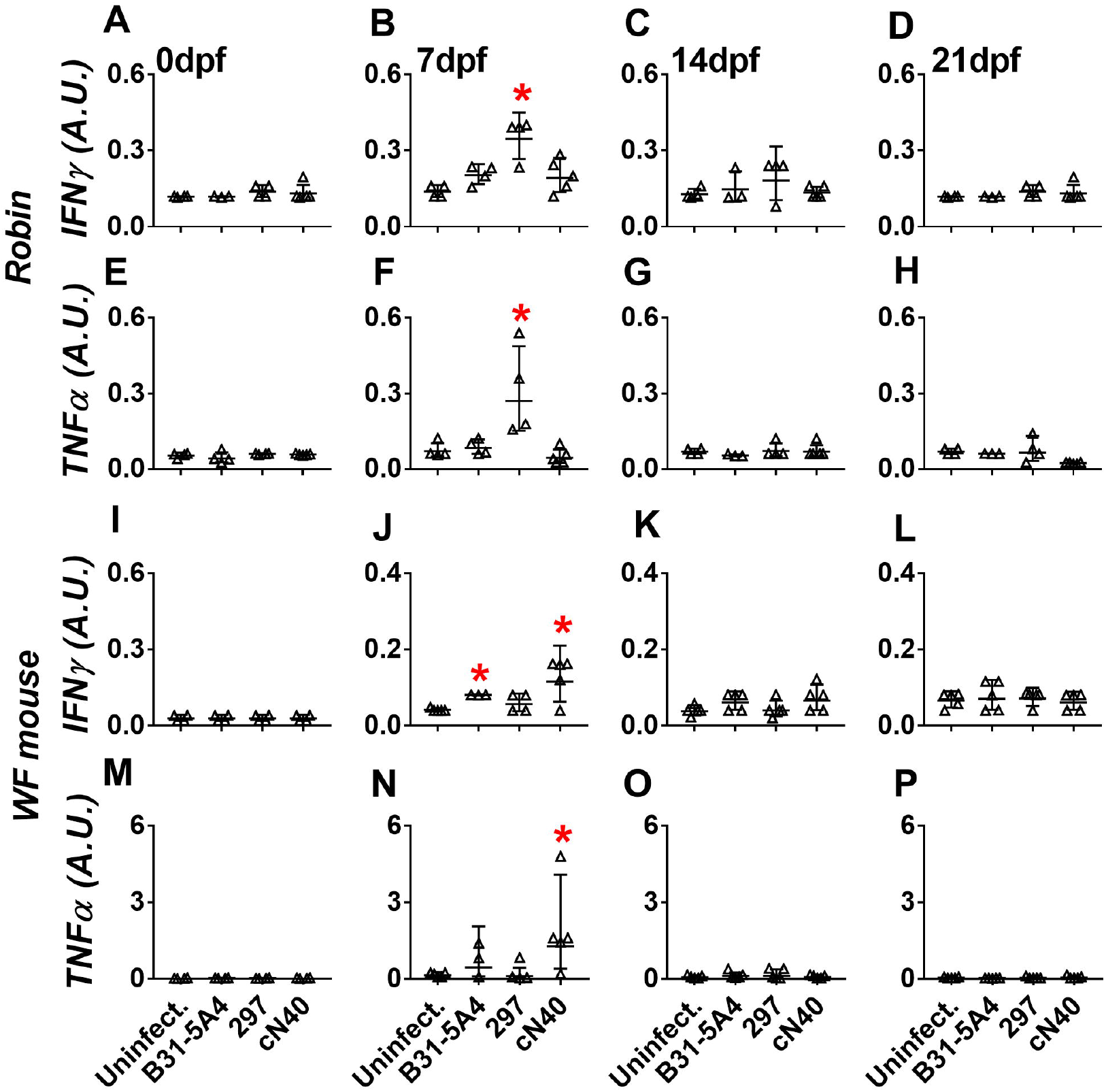
*Borrelia burgdorferi* B31-5A4, 297, and cN40 triggered different levels of pro-inflammatory cytokines at early stages of robin and white-footed mice infection. *Ixodes scapularis* nymphs infected with *B. burgdorferi* B31-5A4, 297, cN40, or naïve nymphs (Uninfect.) were allowed to feed to repletion on **(A to H)** American robins or **(I to P)** white-footed (WF) mice. The sera were obtained **(A, E, I, and M)** prior to tick feeding or at **(B, F, J, and N)** 7-, (**C, G, K, and O)** 14-, or **(D, H, L, and P)** 21-days post nymph feeding (“dpf’). The levels of IFNγ **(A to D and I to L)** and TNFα **(E to H and M to P)** were determined using quantitative ELISA. Shown are the geometric mean ± geometric standard deviation of five white-footed mice per group or robins (3, 4, 5, and 4 for the strains B31-5A4-, 297-, cN40-infected robins or uninfected robins per group, respectively). Significant differences (p < 0.05) in the cytokine levels relative to robins or white-footed mice fed on by naïve nymphs (“*”) are indicated.

### Robins and white-footed mice developed distinct levels of bactericidal antibodies during early infection of *B. burgdorferi* B31-5A4, 297, or cN40

We aimed to quantitatively measure the levels of antibodies induced by the infection of B31-5A4, 297, and cN40. To avoid the confounding factors of strain-specific antibody recognition observed previously (10, 33), we mixed these strains and determined the IgG titers against such mixtures in robins infected with B31-5A4, 297, or cN40. We found that robins infected with any strain develop significantly higher anti-spirochete IgG titers than uninfected robins at 14, 21, and 28dpf whereas only 297-infected robins had significantly higher tiers of IgG than uninfected robins at 7dpf (Fig. 6B to E). cN40 triggered significantly greater levels of IgG antibodies compared to the other strains at 21 and 28 dpf whereas 297 induced IgG titers significantly higher than those titers triggered by other strains at 7dpf (Fig. 6B, D and E). Further, we found that white-footed mice infected with each of these strains had anti-spirochete IgG titers at more robust levels than uninfected mice at 14, 21, and 28dpf whereas only cN40-infected mice developed significantly higher tiers of IgG than uninfected white-footed mice at 7dpf (Fig. 6G to J). cN40 induced significantly lower titers of such IgG at 21 and 28dpf but significantly higher titers at 7dpf, compared to B31-5A4 and 297 (Fig. 6G, I and J). These results indicate that B31-5A4, 297, and cN40 differ in their ability to induce antibodies, and such differences depend on the hosts and infection onset.

**Figure 6.**
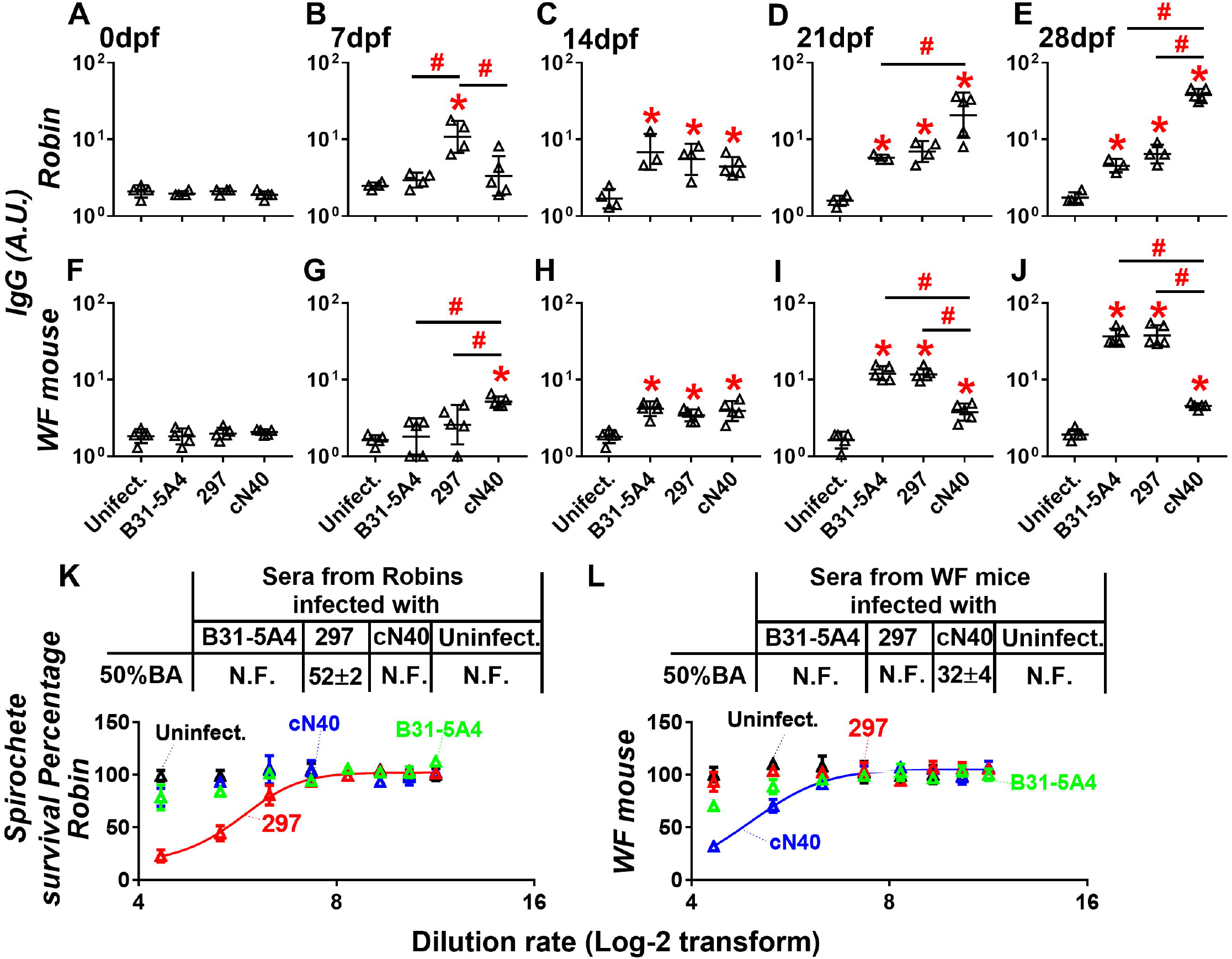
*Borrelia burgdorferi* B31-5A4, 297, and cN40 differed in the ability to induce antibodies against spirochetes in robin and white-footed mice during infection. *Ixodes scapularis* nymphs infected with *B. burgdorferi* B31-5A4, 297, or cN40, or naïve nymphs (Uninfect.) were allowed to feed to repletion on **(A to E)** American robins or **(F to J)** white-footed (WF) mice. The sera were obtained at **(A and F)** 0-, **(B and G)** 7-, **(C and H)** 14-, **(D and I)** 21-, or **(E and J)** 28-days post nymph feeding (“dpi”). The levels of IgG against the mixture of *B. burgdorferi* strains B31-5A4, 297, and cN40 (1×10^6^ spirochetes per strain in each microtiter well) were determined using quantitative ELISA. Shown are the geometric mean ± geometric standard deviation of five white-footed mice or robins per group (3, 4, 5, and 4 for the strains B31-5A4-, 297-, cN40-infected robins or uninfected robins per group, respectively). Significant differences (p < 0.05) in the antibody titers relative to robins or white-footed mice fed on by naive nymphs (“*”) are indicated. Sera from the **(K)** American robins and **(L)** white-footed (WF) mice at 7 days after fed on by *I. scapularis* nymphs carrying *B. burgdorferi* strains B31-5A4, 297, or cN40, or naïve nymphs (Uninfect.) were collected. These sera were serially diluted as indicated and mixed with **(K)** chicken or **(L)** guinea pig complement and the mixture of the strains B31-5A4, 297, and cN40 (1×10^7^ spirochetes per strain in each reaction). After incubation for 24 h, surviving spirochetes were quantified from three fields of view for each sample microscopically. The experiment was performed at three independent occasions. The survival percentage was derived from the proportion of serum-treated to untreated spirochetes. Data shown are the mean ± standard error of the mean (SEM) of the survival percentage from three replicates in one representative experiment. The 50% borreliacidal dilution of each serum sample (50% BA), representing the dilution rate that effectively killed 50% of spirochetes, was obtained from curve-fitting and extrapolation of data shown in the table above respective panels. The 50% BA of some strain-sera pairs could not be determined because no robust killing was observed, resulting in curves that did not fit (N.F.).

The levels of antibodies against B31 −5A4, 297, or cN40 at 7dpf in robins and white-footed mice negatively corresponded to the ability of these hosts to carry spirochetes (Fig. 2B and G). This finding raised a hypothesis that such correspondences are mediated by bactericidal activities of those antibodies. We thus tested this hypothesis by incubating a mixture of B31-5A4, 297, and cN40 with different dilution rates of sera from robins or white-footed mice infected with each of these strains at 7dpf. We found that B31 −5A4- and cN40-derived robin sera did not kill spirochetes at any of the tested dilution rates, similar to sera from uninfected robins. However, 297-derived sera eliminated 50% of spirochetes at the dilution rate of 1:52× (Fig. 6K). Conversely, B31-5A4- and 297-derived white-footed mouse sera did not eliminate spirochetes, similar to the sera from uninfected mice. However, cN40-derived sera eliminated 50% of spirochetes at the dilution rate of 1:32× (Fig. 6L). These finding showed different levels of bactericidal activities from the antibodies produced by B31-5A4-, 297-, or cN40-infected robins and white-footed mice early during infection.

## DISCUSSION

Theory predicts that generalist vectors should select for generalist pathogens, to minimize the latter’s loss to incompetent hosts. Host-specialized genospecies in *B. burgdorferi* s.l. thus represent a paradox, with the intriguing possibility of incipient host specialization within genotypes of the generalist *B. burgdorferi*. We present evidence of molecular mechanisms that differentially influence the ability of three strains of *B. burgdorferi* to colonize, disseminate to distal tissues, evade host immune responses, and be transmitted from the host to feeding ticks between two representative natural reservoir hosts (Fig. 7). We found that cellular and immunological mechanisms act mostly synergistically, resulting in increased fitness of strain cN40 in robins and 297 in white-footed mice. Contrary to theoretical expectations, strain B31-5A4 was able to efficiently infect both hosts, with higher fitness in white-footed mice than the ‘specialized’ strain, 297, and exhibited intermediate fitness in robins. The synergistic nature of those cellular and immunological mechanisms indicates a strong selective pressure for the evolution of host specialization, as predicted by the MNP theory. However, the higher overall fitness of B31-5A4 does not support the existence of tradeoffs, with this strain having high fitness in both representative hosts. This higher fitness advantage is consistent with the high overall prevalence of the genotype of this strain (RST type 1/OspC type A) circulating in Lyme disease prevalent regions of North America (10, 13, 34–37).

**Figure 7.**
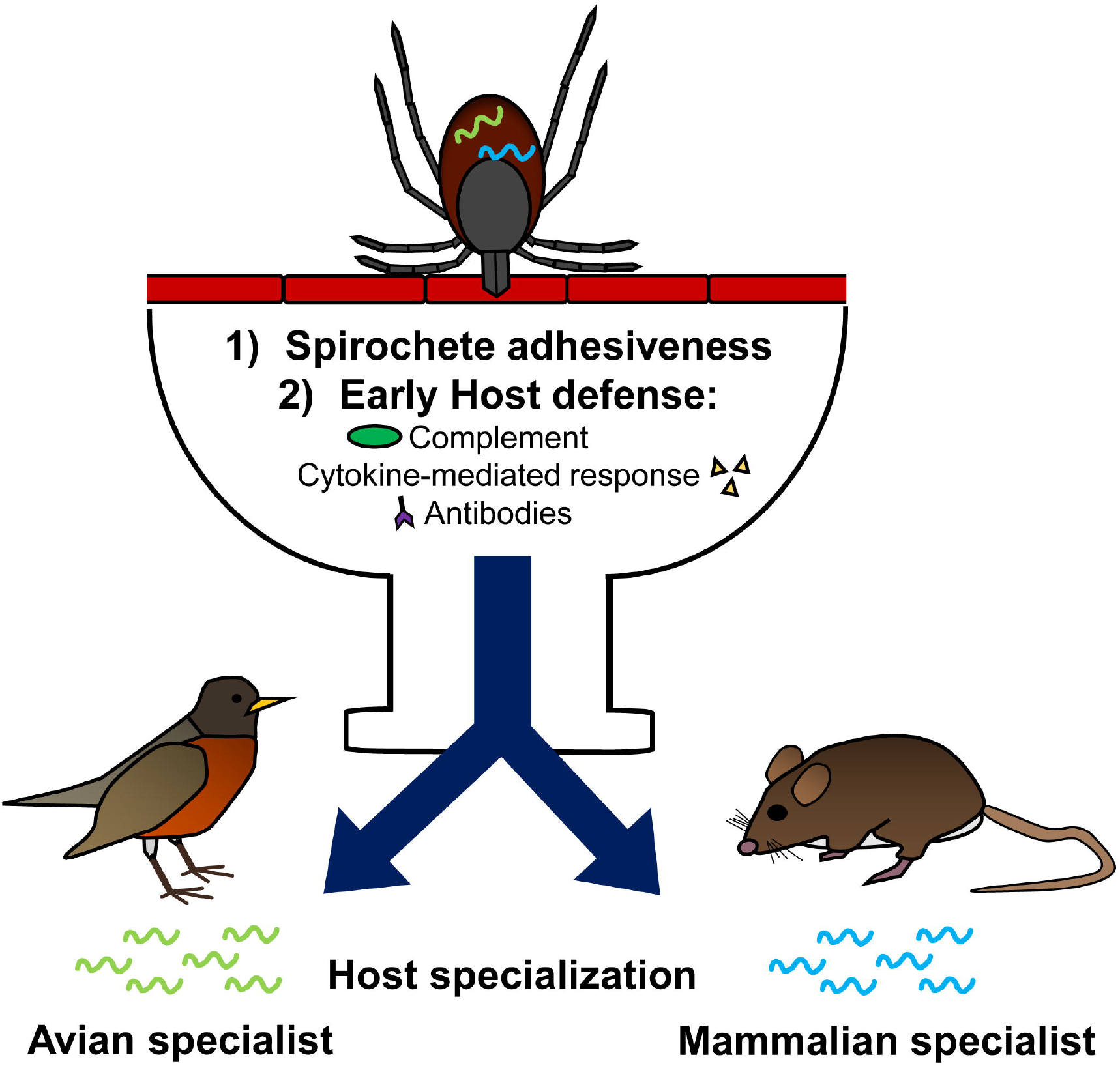
The schematic diagram showing the model supported by this study. Upon *B. burgdorferi* transmission from ticks to hosts, multiple cellular and immunological mechanisms may play in concert to determine the fitness of spirochetes in hosts. These mechanisms include the ability of *B. burgdorferi* to attach to cells or tissues (spirochete adhesiveness) and escape from host immune clearance (early host defense), such as complement, cytokine-mediated responses, and antibodies. Both mechanisms applied on spirochetes would lead to host specialization, resulting in the association of *B. burgdorferi* with particular hosts (i.e. mammal or avian specialists).

Multiple mechanisms were tested in this study for their role in contributing to host specialization of *B. burgdorferi*. At one day after intradermal infection of spirochetes, we found that *B. burgdorferi* B31-5A4 and cN40 colonized robin inoculation sites (skin tissue) more efficiently than 297, whereas B31-5A4 and 297 colonized the inoculation sites of white-footed mice at significantly higher levels than cN40. We also found that these strains differed in their capability to attach to robin and white-footed mouse skin fibroblasts *in vitro.* These results showed that strain-to-strain variability of fibroblast adhesion is host-dependent and corresponds to the ability of *B. burgdorferi* spirochetes to colonize the inoculation sites of these hosts immediately after infection. This supports prior findings of Lyme borreliae strains varying in their adhesive activities (38), demonstrating the role of such cellular processes (adhesion) in conferring strain-host associations.

However, the levels of robin fibroblast adhesion of B31-5A4, 297, and cN40 did not perfectly agree to strain colonization of other tissues (e.g. brain and liver) in robins at 64 days after being fed on by *B. burgdorferi*-infected nymphs. Additionally, strain 297 colonized robin skin at similar levels to B31 and cN40 at 64 dpf, but this strain attached to robin fibroblast at lower levels than other strains. These disagreements between fibroblast adhesion/early colonization of inoculation sites and tissue colonization at later stages suggest the possibility of adhesion-independent mechanisms to determine strain-host associations. In fact, the complement system is the first line of immune defense in vertebrate animal sera and confers Lyme borreliae clearance as soon as infection begins (39, 40). Our findings of greater percentages of B31-5A4 and cN40 surviving in robin sera, compared to those of 297, mirror the trends of these spirochete strains being maintained at earlier infection onsets (e.g. 14 dpf) and early bloodstream survival of *B. burgdorferi* (e.g. 7 dpf). These results imply that the host complement plays a role in determining spirochete genotype-specific robin competence. In contrast, all three *B. burgdorferi* strains showed similar levels of survival in the presence of white-footed mouse sera, suggesting a non-complement-mediated defense mechanism to dominate the association of white-footed mice with these spirochetes.

In addition to complement, we compared the cytokine production in responding to the presence of genotypically distinct *B. burgdorferi*. Cytokines are generally triggered shortly after pathogen invasions, often leading to activation of different pathways and downstream bactericidal responses as a bottleneck of infection initiation (i.e. recruitment of innate and adaptive immune cells) (41–45). *Borrelia burgdorferi* strain-to-strain variability in inducing pro-inflammatory cytokines has been observed in humans, *Mus musculus,* and *P. leucopus* mice or from cells derived from these animals (46–52). In humans, the levels of multiple cytokines are positively correlated with disease severity but are independent from spirochete burdens (46, 47, 53–55). We found that the three tested *B. burgdorferi* strains differed in their ability, in robins and white-footed mouse fibroblasts, to induce the expression of two pro-inflammatory cytokines and their related proteins, IFNγ, TNFα, and TNFα-induced proteins. Such differences also matched the variation of these strains to trigger IFNγ and TNFα during early infection. The ability of these strains to induce those pro-inflammatory cytokines were negatively associated with larval percent positivity from the infection of these *B. burgdorferi* strains. These results raised the hypothesis that less robust host cytokine induction by spirochetes facilitates reservoir competence, and differences in cytokine induction by genotypically distinct spirochetes shape Lyme borreliae-reservoir associations.

Following the induction of innate immune responses, the adaptive responses including antibodies are essential in Lyme borreliae clearance by vertebrates (41, 56, 57). Distinct levels of antibodies were observed during the infection time period by different spirochete strains (10, 58). We observed at 7 dpf that 297 and cN40 triggered significantly higher titers of anti-*B. burgdorferi* IgG in robins and white-footed mice, respectively, compared to the other spirochete strains. Furthermore, *B. burgdorferi* was selectively eliminated by the sera collected at 7dpf from robins infected with 297 or white-footed mice infected with cN40, suggesting that the antibody response varies among different spirochete strain-host pairings. Interestingly, a previous study showed variable IgG levels against lp36-and lp28-1-derived proteins in white-footed mice infected with different *B. burgdorferi* strains (10). However, in that study, those lp36- and lp28-1-derived proteins were produced from a single strain of *B. burgdorferi (B. burgdorferi* strain B31), leading to the possibility that allelic-specific recognition is the result of antigenic sequence variation among spirochete strains (10, 33, 59–61). Thus, the possibility of the antibodies against other spirochete proteins to confer *B. burgdorferi* clearance may not be completely ruled out. Our finding of early antibody-mediated bactericidal activities in robin and white-footed mouse sera highlights the potential role of antibodies in dictating pathogen-host associations. We observed high titers of antibody responses at late time points (21 and 28 dpf) in robins infected with cN40 and mice infected with B31-5A4- or 297. Nonetheless, cN40 was persistently maintained in robins whereas B31-5A4 and 297 efficiently survived long-term in white-footed mice. Such observations are consistent with the evidence of pathogen-specific antibodies present in persistently infected animals (62–64). These results thus suggest inefficient spirochete elimination by those antibodies at later stages of infection. That inefficient pathogen killing could be because of the onset-dependent reduction of highly antigenic spirochete proteins (i.e. downregulation of OspC at late infection onsets) (65–69), constantly changing features of antigen sequences to evade antibody responses (i.e. VlsE) (70–72), or the rapid dissemination of spirochetes to the sites less vulnerable to antibody-mediated clearance (73).

Our results suggest spirochete adhesiveness and early immune responses as the cellular and immunological mechanisms that differentially confer spirochete fitness in reservoir-strain-host combinations (Fig. 7). Both mechanisms could play in concert to determine *B. burgdorferi* strain-to-strain variation in host fitness. Our findings support previous studies which determined that some polymorphic Lyme borreliae proteins confer these immune response functions in a variant-specific manner, and such manner is host dependent (30, 59, 62, 63, 74–80). The potential for host specialization driving genome-wide diversification is illustrated by the recent definition of a new species of *B. burgdorferi* s.l., *B. bavariensis,* which was previously considered a genotype of the avian associated species, *B. garinii.* Such a redefinition is based on the variation in chromosomal housekeeping genes and the association of this species with rodents, unlike other *B. garinii* strains (81). This reclassification reflects how host adaptation can lead to speciation in Lyme borreliae (82). Prior field studies along with our current laboratory evidence showing incipient host specialization raises the possibility of diversification of *B. burgdorferi* in North America into multiple genospecies. Here, we used a controlled laboratory setup to demonstrate differences in fitness of *B. burgdorferi* strains in North American reservoir hosts. The molecular mechanisms supported by our findings provide a potential model of how specific adaptations may lead to specialization even with a significant cost in lost propagules to incompetent hosts. The identified mechanisms can guide future empirical and modeling studies to understand the role of hostpathogen-vector interactions in shaping the microparasite host ranges and their potential for pathogen spillover into livestock, wildlife or humans.

## MATERIALS AND METHODS

### Ethics statement

All experiments involving American robins (*Turdus migratorius*) and white-footed mice *(Peromyscus leucopus)* were performed in strict accordance with all provisions of the Animal Welfare Act, the Guide for the Care and Use of Laboratory Animals, and the PHS Policy on Humane Care and Use of Laboratory Animals. Additionally, the protocol was approved by the Institutional Animal Care and Use Committee (IACUC) of Wadsworth Center, New York State Department of Health (Protocol docket number 18-412 and 19-451), and Columbia University (Protocol number AC-AAAY2450), and the City University in New York Advanced Science Research Center (Protocol number ASRC AUP 2016-20). To mistnet robins, personnel were approved on scientific collecting permits USFWS Collecting Permit MB035731 and NYSDEC Permit #1236.

### Bird, Mouse, Tick, and Bacterial strains

From June-July, 2019 and July, 2020, American robins were mist netted around the property of Griffin Laboratory of Wadsworth Center, New York State Department of Health at Albany, NY. This site was selected because of previously reported low tick abundance, thus minimizing the probability of previous tick exposure. Sera were collected from 32 and 17 robins in 2019 and 2020, respectively, using BD Microtainer Capillary Blood Collector tubes (Fisher Scientific, Hampton, NH, USA) to assess previous infection with *B. burgdorferi* infection by the methods previously described (31). To confirm seronegative status, robins were quarantined for two weeks at 25°C on a 12L:12D (light: dark) cycle by housing them in cages with a wire bottom, under which a water moat was placed. If replete ticks were found while the birds were in quarantine, quantitative PCR (qPCR) was used to determine the spirochete burdens in those ticks (see “Quantification of *B. burgdorferi* in infected ticks, tissues, and blood samples” for more details). The birds with spirochete positive ticks attached were removed from the experiments. After two weeks of quarantine, robins were subjected to additional serological examination as described above, and 40 seronegative juvenile (hatch year) robins were considered non-infectious and used in this study.

White-footed mice were purchased from the *Peromyscus* Genetic Stock Center at the University of South Carolina (Columbia, SC). Non-sibling mice were bred in-house at Columbia University. Immunodeficient Fox Chase SCID mice (C.B.17 SCID) were obtained from Charles River (Boston, MA) and used to generate infected nymphs for each *B. burgdorferi* strain as described in the “Mouse infection experiments by ticks” section. *Ixodes scapularis* larvae were purchased from the National Tick Research and Education Center, Oklahoma State University (Stillwater, OK). Mice and birds were housed individually and maintained at 21 to 24°C on a 14L:10D (light: dark) cycle and handled humanely. The *B. burgdorferi* strains used in this study were cultivated in BSK-II completed medium as described in Table 1 (83).

### Determination of spirochete growth curves and generation time

*Borrelia burgdorferi* strains B31-5A4, 297, and cN40 were cultivated in BSK-II complete media at 33°C in the initial concentration of 5×10^6^ ml^-1^. The concentration of spirochetes was measured prior to incubation and at 24-, 48-, 72-, 96-, 120-, 144-, and 168-h post incubation using a Nikon Eclipse E600 dark field microscope (Nikon, Melville, NY). The generation time of each spirochete strain at the exponential phase was calculated as described previously (84).

### Robin, C.B.17 SCID mouse, and white-footed mouse infection by intradermal inoculation

Four to eight week old male or female white-footed mice, American robins, or C.B.17 SCID mice were intradermally inoculated, using a 27-gauge needle, with *B. burgdorferi* B31-5A4, 297, cN40, or BSK-II medium without rabbit sera (1×10^5^ bacteria per C.B.17 SCID mouse or 1 ×10^4^ bacteria per robin or white-footed mouse) as a control (63). The plasmid profiles and the presence of the shuttle plasmids of each of these *B. burgdorferi* strains were verified prior to infection to ensure no loss of plasmids, as described previously (85–87). Both robins and white-footed mice were euthanized at one day post injection. The inoculation site of the skin and blood from robins and white-footed mice were collected to quantitatively evaluate spirochete burdens as described in the “Quantification of *B. burgdorferi* in infected ticks, tissues and blood samples” section.

### Quantification of *B. burgdorferi* in infected ticks, tissues, and blood samples

The white-footed mouse- or robin-derived replete nymphs were mixed with glass beads and homogenized by a Precellys 24 High-Powered Bead Mill Homogenizer (Bertin, Rockville, MD). DNA was extracted from blood and tissue samples of white-footed mice, robins and homogenized ticks using an EZ-10 Genomic DNA kit (Biobasic, Amherst, NY). The quantity and quality of DNA for each tissue sample was assessed by measuring the concentration of DNA and the ratio of the UV absorption at 280 to 260 using a nanodrop 1000 UV/Vis spectrophotometer (ThermoFisher Scientific, Waltham, MA). The amount of DNA used in this study was 100 ng for each sample, and the 280:260 ratio was between 1.75 to 1.85, indicating the lack of contamination by RNA or proteins. qPCR was performed to quantitate bacterial loads. *Borrelia burgdorferi* genomic equivalents were calculated using an Applied Biosystems 7500 Real-Time PCR system (ThermoFisher Scientific) in conjunction with PowerUp SYBR Green Master Mix (ThermoFisher Scientific), based on amplification of the *B. burgdorferi 16S rRNA* gene using primers 16S rRNAfp and 16S rRNArp (Table S2), as described previously (88). Cycling parameters for SYBR green-based reactions were 50°C for 2 min, 95°C for 10 min, 45 cycles of 95°C for 15 s, and 60°C for 1 min. The number of *16S rRNA* copies was calculated by establishing a threshold cycle (Ct) standard curve of a known number of *16S rRNA* gene copies extracted from *B. burgdorferi* strain B31, and compared to the Ct values of the experimental samples. To ensure low signals were not simply a function of the presence of PCR inhibitors in the DNA preparation, five samples of blood, tibiofemoral joints, bladders of white-footed mice and the skin, brain, and heart of robins in the B31-5A4 experimental group were applied to qPCR to determine the levels of β-actin from white-footed mice (pActfp and pActrp) and robins (rActfp and rActrp), respectively (Table S2) (63, 89). Note that the primers used to determine the expression of the genes encoding robin actin were based on the mRNA sequences that were translated to protein (Genbank Accession number: PYHW01009720.1) from a closely related avian host, rufous-bellied thrush *(Turdus rufiventris)* due to the lack of sequence information of American robin actin. As predicted, we detected 10^7^ copies of the actin gene from 100ng of each DNA sample in robins and white-footed mice, ruling out the presence of PCR inhibitors in these samples.

### Isolation of robin fibroblasts

The procedures to isolate robin fibroblasts were described previously (90). Five 5mm × 5mm sections of skin were removed from the breast of euthanized robins, washed twice in PBS buffer, and then incubated in a transferring solution until the next step was ready to be performed. The constituents of the transferring solution included Dulbecco’s modified Eagle medium (DMEM) (Wadsworth media & tissue culture core) with glucose (4.5mg ml^-1^) (Sigma-Aldrich, St. Louis, MO), sodium pyruvate (110mg l^-1^) (Sigma-Aldrich), L-glutamine (ThermoFisher Scientific), supplemented with 10% heat-inactivated fetal bovine serum (ThermoFisher Scientific), 2% heat-inactivated chicken serum (Biowest, Riverside, MO, USA), and antibiotics (100U ml^-1^ of mixture of penicillin and streptomycin) (ThermoFisher Scientific). The skin was then submerged in 70% ethanol (Sigma-Aldrich) for 30s, minced using sterile scalpels, and placed in collagenase B at 37°C overnight (ThermoFisher Scientific). We then applied the mixture of the cells to a 20μm cell strainer and spun down the samples. The pellets containing fibroblasts were re-suspended in growth media and incubated at 37°C with 5% CO2. Growth media had the same components as the transferring solution except it was supplemented with amphotericin B (ThermoFisher Scientific) to a concentration of 0.25μg/ml. The cells were harvested by trypsinization using Trypsin (0.25% trypsin in DMEM media, ThermoFisher Scientific) and used in the spirochete attachment experiment.

### Determination of the levels of spirochete attachment to robin and white-footed mouse fibroblasts

The primary fibroblasts from the neck skin of American robins (see the previous section) and ears of white-footed mice were acquired commercially (#AG22353, Coriell Institute for Medical Research, Camden, NJ) and cultivated on cover slips in 24-well plates (2 × 10^5^ cells per well). *Borrelia burgdorferi* strains B31-5A4, 297, cN40, or B314 were suspended in BSK-II medium and added to prepared plates (2 × 10^6^ spirochetes per well). The plates were centrifuged at 106 g for 5 min and then rocked at room temperature for 1 h. After removing unbound bacteria through washing each well with PBS containing 0.2% BSA, the bound bacteria and cells on the cover slips were fixed using 100% chilled methanol for 1 h followed by blocking with PBS containing 0.2% BSA for 1 h. After washing with PBS containing 0.2% BSA, the cover slips were incubated with a fluorescein isothiocyanate (FITC)-conjugated goat anti-*B. burgdorferi* polyclonal antibody (Abcam, Cambridge, MA) for 1 h and mounted using ProLong Gold antifade mountant with DAPI fluorescent stain (ThermoFisher Scientific). The spirochetes (green) and the DNA from spirochetes and the nuclei of fibroblasts (blue) were then visualized under overlaid FITC and DAPI filters using an Olympus BX51 fluorescence microscope (Olympus Corporation, Waltham, MA) (Fig. S3). The number of spirochetes from three fields of view were counted in four independent events. The results were presented as the average number of spirochetes per 50 fibroblast cells.

### Determination of the relative expression levels of the genes encoding IFNγ and TNFα or TNFα-induced proteins in fibroblasts by quantitative reverse transcription PCR (RT-qPCR)

The procedures for the examination of expression levels in the genes encoding cytokines of fibroblasts were described previously (52). In brief, the primary fibroblasts from robins and whitefooted mice were cultivated in 24-well plates (2 × 10^5^ cell per well). When the cells were greater than 80% confluent, *B. burgdorferi* strains B31-5A4, 297, or cN40 suspended in BSK-II medium were added to corresponding wells on each plate (2 × 10^6^ and 2 ×10^7^ spirochetes per well for the cell to spirochete ratio (MOI) of 1:10 and 1:100, respectively) for 24h. The cells incubated with BSK-II medium without bacteria (mock-treated cells) were included as a control.

After incubation, the supernatant was removed, and the cells were washed with PBS buffer. These fibroblasts were then suspended in Trizol (ThermoFisher Scientific) at room temperature for 1 h to inactivate RNase. The procedure of RNA extraction was performed using Direct-Zol RNA MiniPrep Plus Kit (Zymo Research, Irvine, CA) as previously described (84), and DNA was removed using RQ1 RNase-Free DNase (Promega, Madison, WI). We then synthesized cDNA using these RNA samples (1 μg per sample) by qScript cDNA SuperMix (Quanta Bioscience, Beverly, MA). The expression levels of the house keeping genes encoding the 18S rRNA gene from robins (Genbank Accession number: M59402.1) or white-footed mice (Genbank Accession number: AY591913.1) were included as controls. The primers used to quantitate the expression of the genes encoding white-footed mouse IFNγ, TNF, and 18S rRNA, and robin IFNγ, TNFα- induced proteins, and 18S rRNA are listed in Table S2 (89). Note, because of the lack of sequence information for robin cytokines, the primers to determine the expression of these cytokines were based on the mRNA sequences of IFNγ (Genbank Accession number: PYHW01010552.1) and TNFα-induced proteins (Genbank Accession number: PYHW01009717.1) from rufous-bellied thrush (*Turdus rufiventris*). The quantity and quality of cDNA for each sample was evaluated by obtaining the concentration of DNA and the ratio of the UV absorption at 260 and 280 using a Nanodrop 1000 UV/Vis spectrophotometer (ThermoFisher, Waltham, MA). The resulting ratio between 1.75 and 1.85, indicated the lack of RNA or protein contamination. Samples were applied to an Applied Biosystems 7500 Real-Time PCR System (ThermoFisher) in conjunction with PowerUp SYBR Green Master Mix (ThermoFisher) to detect the expression levels of the abovementioned genes. The cycling parameters were 50°C for 2 min, 95°C for 10 min, and 45 cycles of 95°C for 15 s, and 49°C for 1 min, and the resulting values of threshold cycles (Ct) were determined from duplicate experiments in three independent events. The relative expression of the genes encoding IFNγ, TNF, TNFα-induced proteins, or 18S rRNA was presented by normalizing the Ct-derived from each of these cytokines to that of β-actin from respective animals through the following equation (Equation 1).

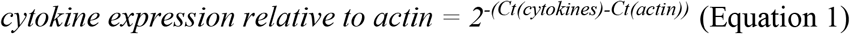

### Generation of *B. burgdorferi-infected* nymphal ticks

Four-week-old male and female C.B.17 SCID mice were injected with a concentration of 1×10^5^ of either *B. burgdorferi* strain B31-5A4, 297, or cN40 via subcutaneous injection. The plasmid profiles and the presence of the shuttle vector of each of these *B. burgdorferi* strains were verified prior to injection to ensure no loss of plasmids, as previously described (85–87). SCID mice inoculated with BSK-II media without rabbit sera were used to generate uninfected nymphs. A 3 mm ear biopsy was collected 7 days post injection from each mouse inoculated with a *B. burgdorferi* strain and DNA was extracted using a DNeasy Blood and Tissue kit (Qiagen, Germantown, MD). DNA from ear tissue was subjected to qPCR analysis to verify infection in each host (30). At 14 days post injection, approximately 200 larvae were placed in the ears of anesthetized mice using a paint brush. Mice were anesthetized with isoflurane for 45 min to allow larvae to attach before being housed individually in water bath cages, larvae were allowed to feed to repletion, as described previously (30, 91). Engorged larvae were collected, cleaned with 10% bleach and 70% ethanol solutions, and stored in an incubator at 21°C, 95% relative humidity, and a 14L:10D light: dark cycle. Larvae molted into the nymphal life stage in approximately 4-6 weeks and were used in the subsequent experiments with white-footed mice and robins.

### Robin and white-footed mouse infection by nymphs

The timeline of experimental procedures is provided in Fig. 2A. Basically, unfed nymphs carrying B31-5A4, 297, cN40 or unfed, uninfected nymphs were placed in the ear canals of each mouse (*n* = 5 mice per group, 5 nymphs in each ear). Mice were maintained under anesthesia for 60 min to allow nymphs to attach before being placed individually in water bath cages. Approximately 100 to 150 xenodiagnostic larvae were placed on each mouse at 14, 28, 35, and 56 days post nymph feeding using the aforementioned procedure. To permit flat nymphs to feed on robins, the birds were placed into a PVC pipe (~2.5-3.0 inches in diameter; 10 inches in length) as described (92). After unfed nymphs carrying B31-5A4, 297, cN40 or unfed, uninfected nymphs were placed on these robins (*n* = 5 robins per group, 10 nymphs per bird), the PVC pipes were covered by a fine mosquito net (‘no-see-um’ netting, Skeeta, Bradenton, FL) secured with a rubber band. The birds were then moved into wire bottom cages with a water moat. These tick-infested robins were kept in a dark room for 1 h to minimize grooming and allow ticks to attach before they were released and returned to their cages. To allow xenodiagnostic larvae to feed on robins, approximately 100 to 200 naïve larvae were placed on robins restrained in the PVC pipes as described above at 14, 28, 35, and 56 days post nymph feeding.

Blood samples were collected from each animal (robins and white-footed mice) prior to nymph feeding and at 7, 14, 21, and 28 days post nymph feeding. The skin, heart, liver, and brain from robins and the ears, tibiofemoral joints, heart, and bladder from white-footed mice were obtained at 64 days post nymphs feeding. The blood, tissues, and replete nymphs collected from each animal were analyzed via qPCR to determine spirochete burdens as described in the section “Quantification of *B. burgdorferi* in infected ticks, tissues, and blood samples”.

### Serum resistance assays

Serum resistance of *B. burgdorferi* strains was determined as described previously with modifications (30, 84). The mid-log phase *of B. burgdorferi* strains B31-5A4, 297, and cN40 as well as a high passage, serum sensitive strain B313 (negative control) were cultivated in triplicate. The resulting spirochete culture was diluted to a final concentration of 5×10^6^ bacteria per milliliter into BSK-II medium without rabbit sera. The cell suspensions were mixed with sera collected from naïve white-footed mice or robins (60% spirochetes and 40% sera) in the presence or absence of 2 μM of cobra venom factor (CVF) or recombinant *Ornithordorus moubata* complement inhibitor (OmCI). The generation of recombinant OmCI has been described previously (31). The sera were incubated at 65°C for 2 h (heat-inactivated sera) and included as a control. At 0 and 4 h following incubation with sera, the number of motile spirochetes was measured under dark field microscopy. The percent survival of *B. burgdorferi* was calculated by the normalization of motile spirochetes at 4 h post incubation to that immediately after incubation with sera.

### Determination of the relative levels of IFNγ and TNFα in sera by ELISA

Due to the lack of commercially available ELISA kits to detect robin cytokines, the kits for chicken IFNγ (ThermoFisher) and TNFα (Genorise, Glen Mills, PA) were used to measure the levels of those cytokines in robins based on the absorption values derived from ELISA. Similarly, the ELISA kits to determine the levels of IFNγ and TNFα from house mouse *(Mus muscuslus)* (Tonbo Bioscience, San Diego, CA) were utilized to detect those cytokines in white-footed mice. We observed the antibodies in the ELISA kits cross reacted with IFNγ and TNFα from robins and white-footed mice. However, the recombinant proteins provided in these kits to generate standard curves were based on the sequences from chicken and house mouse IFNγ and TNFα, potentially leading to inaccuracy of quantification for those cytokines by normalizing the results obtained from robin and white-footed mouse samples to standard curves. Therefore, we chose to present reservoir animal cytokines as relative levels (arbitrary unit; A.U.), similar to a previous study (93). In brief, the monoclonal capture antibodies that recognized chicken or house mouse IFNγ or TNFα were coated on microtiter plate wells. After blocked by 5% PBS-BSA, different dilutions (1:100×, 1:300×, or 1:900×) of sera from robins and white-footed mice at 0, 7, 14, or 21 days post nymph feeding were added to the wells. Wells were washed using PBST buffer (PBS with 0.5% of Tween 20) and the detection of antibodies against chicken or house mouse IFNγ or TNFα were incubated for 1 h at room temperature. The wells were then washed with PBST buffer and subsequently mixed with a tetramethyl benzidine solution (ThermoFisher). The absorbance level was detected at 620nm for 10 cycles of 60 s kinetic intervals with 10 s of shaking duration in a Sunrise absorbance ELISA plate reader (Tecan, Männedorf, Switzerland). We then obtained the greatest maximum slope of optical density/min per sample multiplied by respective serum dilution factors to represent the relative levels of cytokines shown as arbitrary units.

### Determination of the titers of the antibodies against spirochetes

The titers of IgG against spirochetes were measured as described previously with modifications (94). Basically, microtiter plate wells were coated with a mixture of *B. burgdorferi* B31, 297, and cN40 (1 × 10^6^ spirochetes per strain in a well). After blocking with 5% PBS-BSA, the sera from white-footed mice or robins collected at 7, 14, 21, 28, and 56 days post nymph feeding were diluted in 50μl of PBS (1:100×, 1:300×, or 1:900×) and then added to the wells. After the samples were washed with PBST buffer, the microtiter wells were incubated with antibodies that recognize the Fc region of IgG from *P. leucopus* (1:1,000×, Serocare, Inc, Milford, MA), or wild bird (1:10,000×, Bethyl laboratory, Montgomery, TX). After the addition of these antibodies, tetramethyl benzidine solution (ThermoFisher) was added, and the absorbance was detected at 620nm for 10 cycles of 60 s kinetic intervals with 10 s of shaking duration in a Sunrise absorbance ELISA plate reader (Tecan, Männedorf, Switzerland). The values of greatest maximum slopes of optical density/min/sample were multiplied by respective serum dilution factors as shown as arbitrary units, representing antibody titers.

### Borreliacidal assays

Sera collected from robins and white-footed mice at 7 days post tick feeding were used to measure the bactericidal activity against *B. burgdorferi* as described previously (95). The complement in these serum samples was heat-inactivated by incubating these samples at 55°C for 2 h. The heat inactivated sera were serially diluted in BSK-II media without rabbit sera followed by being mixed with complement-preserved sera from guinea pig (Sigma-Aldrich) or chicken (Biowest, Riverside). Heat-inactivated guinea pig and chicken sera were included as controls. *Borrelia burgdorferi* strains B31-5A4, 297, or cN40 (1 × 10^7^ spirochetes per strain) were then mixed prior to their addition to the reaction. At 24 h after incubation, the surviving spirochetes were quantified by counting the motile bacteria under dark-field microscopy and presented as the proportion of serum-treated to untreated spirochetes. We also calculated the 50% borreliacidal titer (BA), which represents the serum dilution rate that eradicates 50% of spirochetes, using doseresponse stimulation fitting in GraphPad Prism 7.

### Statistical analyses

Because the data were not normally distributed, we used a Kruskal-Wallis test followed by a two-stage step-up method with a Benjamini, Krieger, and Yekutieli correction for all comparisons (96). Geometric means of duplicated individual samples (qPCR) were used in our calculations. Spirochete burdens among the three *B. burgdorferi* strains (B31-5A4, 297, and cN40) and uninfected samples in robins, white-footed mice, or ticks were compared using normalized qPCR quantity values for each individual. Cell adhesion variation among the *B. burgdorferi* strains were analyzed using the number of spirochetes per 50 fibroblast cells observed under fluorescent microscopy while the variation in expression levels of the genes encoding IFN-γ, TNF, or TNFα-induced protein were first normalized by the gene encoding actin and then significant differences between the constitutively expressed gene, actin, and 18S rRNA were determined using RT-qPCR quantity values. However, when comparing the differences between levels of pro-inflammatory cytokines at early stages of host infection, we used quantitative ELISA values. Spirochete survival in treated and untreated host sera was assessed using the number of mobile spirochetes at 4 h post incubation. We used quantitative ELISA values to determine the significant differences among the *B. burgdorferi* strains and levels of IgG antibody production in robins and white-footed mice. All analyses and figure generation were completed using GraphPad Prism 7 software.

## ACKNOWLEDGEMENTS

We thank Thomas Hart and Tristan Nowak for graph editing, and Jen Owen and Jean Tsao for technical support in the protocol of tick infection in robins. We also thank Thomas Hart, Patricia Lederman, Tristan Nowak, and Frank Blaisdell for the husbandry work to maintain tick colony and robins. We are grateful for John Leong and George Chaconas for sharing the *B. burgdorferi* strains B31-5A4, 297, and cN40, Sudha Chaturvedi allowing us to use her Bead Homogenizer, Susan Madison-Antenucci allowing us to use her fluorescence microscope, and Noel Espina for the assistance with that microscope. We thank Wadsworth Media & Tissue Culture Core for preparation of *Borrelia* and American robin and white-footed mouse fibroblast culture medium. This work was supported by NSF IOS-1755370 (DMT, MC and MDW), NSF-IOS1755286 (YL, ADII, ALM, JS, and LDK), New York State Department of Health Wadsworth Center Start-Up Grant (ALM and YL). The funders had no role in study design, data collection and analysis, decision to publish, or preparation of the manuscript. The authors declare that the research was conducted in the absence of any commercial or financial relationships that could be construed as a potential conflict of interest.

## Notes

### Competing Interest Statement

The authors have declared no competing interest.

