## Supplemental Text for "Host specialization in microparasites transmitted by generalist vectors: insights into the cellular and immunological mechanisms"

### SUPPLEMENTARY INFORMATION

### SUPPLEMENTARY FIGURE LEGENDS

**Figure S1. *Borrelia burgdorferi* B31-5A4, 297, and cN40 did not differ in their generation time when cultivated *in vitro*.** $5\times10^6$ cells of *B. burgdorferi* B31-5A4, 297, or cN40 were cultivated in fresh BSK-II medium at 33ºC. The concentration of each of these strains was quantified microscopically at 0-, 24-, 48-, 72-, 96-, 120-, 148-, and 172-h. The assays were performed on three independent occasions; within each experiment, the number of spirochetes was the average of counting in three fields. Shown are the mean number of spirochetes ± SEM of three different experiments. The generation time (G) was calculated and shown in the inlet figure. No significant differences of generation ($P < 0.05$) were observed among these spirochete strains.

**Figure S2. *Borrelia burgdorferi* B31-5A4, 297, and cN40 were not detectable in the bloodstream of robins and white-footed mice at 24 h post infection. (A)** American robins and **(B)** white-footed (WF) mice were intradermally inoculated with $10^4$ spirochetes of *B. burgdorferi* strains B31-5A4, 297, or cN40, or with BSK-II medium without rabbit sera as mock infection (“Mock”). The blood from these animals were collected at 1-day post infection (1dpi) to determine bacterial burdens by qPCR. The bacterial loads in the blood were normalized to 100 ng total DNA. Shown are the geometric mean ± geometric standard deviation of bacterial burdens in those tissues from four robins or five white-footed mice per group. Significant differences ($P < 0.05$) in the spirochete burdens relative to the mock infected group are indicated (‘‘*’’).

**Figure S3. *Borrelia burgdorferi* B31-5A4, 297, and cN40 varied in their ability to attach to fibroblasts derived from robins and white-footed mice.** *Borrelia burgdorferi* strains B31-5A4,

297, cN40, or B314 (negative control) ($2 \times 10^6$ spirochetes) were incubated with fibroblasts ($2 \times 10^5$ cells) from **(A to D)** American robins and **(E to H)** white-footed (WF) mice for 1 h. We mixed those cells with FITC-conjugated goat anti-*B. burgdorferi* polyclonal antibodies, and DAPI was then incubated with these cells to localize the nuclei of those fibroblasts. After fixation, the resulting slides were visualized for the spirochetes under 40-fold magnification (bar, 25 μm). The levels of spirochete attachment were evaluated by counting the number of bacteria per 50 cells under fluorescence microscopy reported in Fig. 1C and D.

**Figure S4. Robin and white-footed mouse fibroblasts treated with *B. burgdorferi* B31-5A4, 297, or cN40 resulted in indistinguishable levels of pro-inflammatory cytokines from mock-treated cells at a spirochete to cell ratio as 1 to 10.** Fibroblasts ($2 \times 10^5$ cells) from **(A to C)** American robins and **(D to F)** white-footed (WF) mice were incubated for 24 h with *B. burgdorferi* strains B31-5A4, 297, or cN40 ($2 \times 10^6$ of spirochetes for spirochete to cell ratio (MOI) at 1:10). Cell media-treated fibroblasts were included as control ("Mock"). After the RNA was extracted from these cells, the expression levels of the gene encoding IFN-γ, TNF, or TNFα-induced protein and the constitutively expressed genes, actin and 18S rRNA from robins and white-footed mice were determined using quantitative reverse transcription polymerase chain reaction. The expression levels of **(A and D)** 18S rRNA, **(B and E)** IFNγ, **(C)** TNFα-induced protein, and **(F)** TNF are presented by normalizing to the expression levels of the gene encoding actin. Each bar represents the mean of three independent determinations ± SD. There was no statistic difference of the relative expression for each of these genes between cells under mock treatment and the cells treated with each of these spirochete strains ($P > 0.05$).

**Figure S5.** ***Borrelia burgdorferi*** **B31-5A4, 297, and cN40 did not vary in their ability to survive in nymphs feeding on robins and white-footed mice.** *Ixodes scapularis* nymphs infected with *B. burgdorferi* strains B31-5A4, 297, or cN40, or naïve nymphs (Uninfect.) were allowed to feed to repletion on American robins or white-footed (WF) mice. The spirochete burdens in the **(A)** flat or replete ticks feeding on **(B)** robins or **(C)** white-footed mice were determined by qPCR. Shown are the geometric mean ± geometric standard deviation of 6 nymphs (for flat naïve nymphs), 6, 5, or 5 nymphs (for flat nymphs carrying the strains B31-5A4, 297, and cN40, respectively), 30 nymphs (for naïve nymphs feeding on mice), or 9, 19, and 41 nymphs (for nymphs carrying the strains B31-5A4, 297, and cN40 feeding on mice, respectively), or 6 nymphs (for naïve nymphs feeding on robins), or 6, 9, and 7 nymphs (for nymphs carrying the strains B31-5A4, 297, and cN40 feeding on robins, respectively). Significant differences ($P < 0.05$) in the spirochete burdens relative to the mock infected group are indicated (‘‘*’’).

### SUPPLEMENTARY TABLES

**Table S1. The *B. burgdorferi* strains used in this study.**

| Strains | Genotype | | Prevalence[c] | Characteristic | References of sources |
|---|---|---|---|---|---|
| | OspC[a] | RST[b] | | | |
| B31-5A4 | A | 1 | 14.4 | Low-passage, clone 5A4 of *B. burgdorferi* strain B31 isolated from a *I. scapularis* tick in Shelter Island, NY, USA | (85, 97) |
| 297 | K | 2 | 19.4 | Low-passage, clone A11/B11 of *B. burgdorferi* strain 297 isolated from human cerebrospinal fluid in MA, USA | (98, 99) |
| cN40 | E | 3 | 4.0 | Low-passage, clone cN40 of *B. burgdorferi* strain N40 isolated from *I. scapularis* ticks in NY, USA. | (100, 101) |
| B313 | A | 1 | | High-passage *B. burgdorferi* B31 missing lp5, lp17, lp21, lp25, lp28-1, lp28-2, lp28-3, lp28-4, lp36, lp38, lp54, lp56, cp9, cp32-4, cp32-6, cp32-8, cp32-9 | (102) |
| B314 | A | 1 | | High-passage *B. burgdorferi* B31 missing lp5, lp16, lp17, lp21, lp25, lp28-1, lp28-2, lp28-3, lp28-4, lp36, lp29, lp38, lp49, lp54, lp56, cp9, cp32-6, cp32-7, cp32-9. | (102) |

[a]See the reference (103, 104)
[b]See the reference (105)
[c]The percent prevalence of the *ospC* genotypes in northeastern USA of the indicated strains (106)

**Table S2. Primers used in this study.**

| Primer/Vector | Sequence* | Amplified DNA fragment |
|---|---|---|
| 16SrRNAfp | gcttcgcttgtagatgagtctgc | *B. burgdorferi 16S rRNA* |
| 16SrRNArp | ttccagtgtgaccgttcacc | |
| pActfp | gcaagcaggagtacgatgag | white-footed mouse *β-actin* |
| pActrp | ccatgccaatgttgtctt | |
| rActfp | tgtagccatccaggctgtgct | Robin *actin* |
| rActrp | gtcacgcacgatttccctct | |
| p18SrRNAfp | tggttcctttggtcgctcgctcc | white-footed mouse *18S rRNA* |
| p18SrRNArp | gaccgggttggttttgatctgat | |
| pIfnarfp | ctggagaccactcggataaatg | white-footed mouse *IFNγ* |
| pIfnarrp | ctcgtacccggagaaagaaag | |
| pTnfsffp | aatgccctcctggccaacg | white-footed mouse *TNF* |
| pTnfsfrp | tcagacccctcgggggtttc | |
| r18SrRNAfp | cctcgatgctcttagctgagtgt | Robin *18S rRNA* |
| r18SrRNArp | cggagtcctattccattattcct | |
| rIfngfp | acagagaagctgtcaagctgg | Robin *IFNγ* |
| rIfngrp | agatttgtcaggtccttgaga | |
| rTnfipfp | agatgctgagtctcatctgg | Robin *TNFα-induced protien* |
| rTnfiprp | catctgctcagtgaatttttg | |
