## Supplementary figures and images for "Host specialization in microparasites transmitted by generalist vectors: insights into the cellular and immunological mechanisms"

### Figure S1

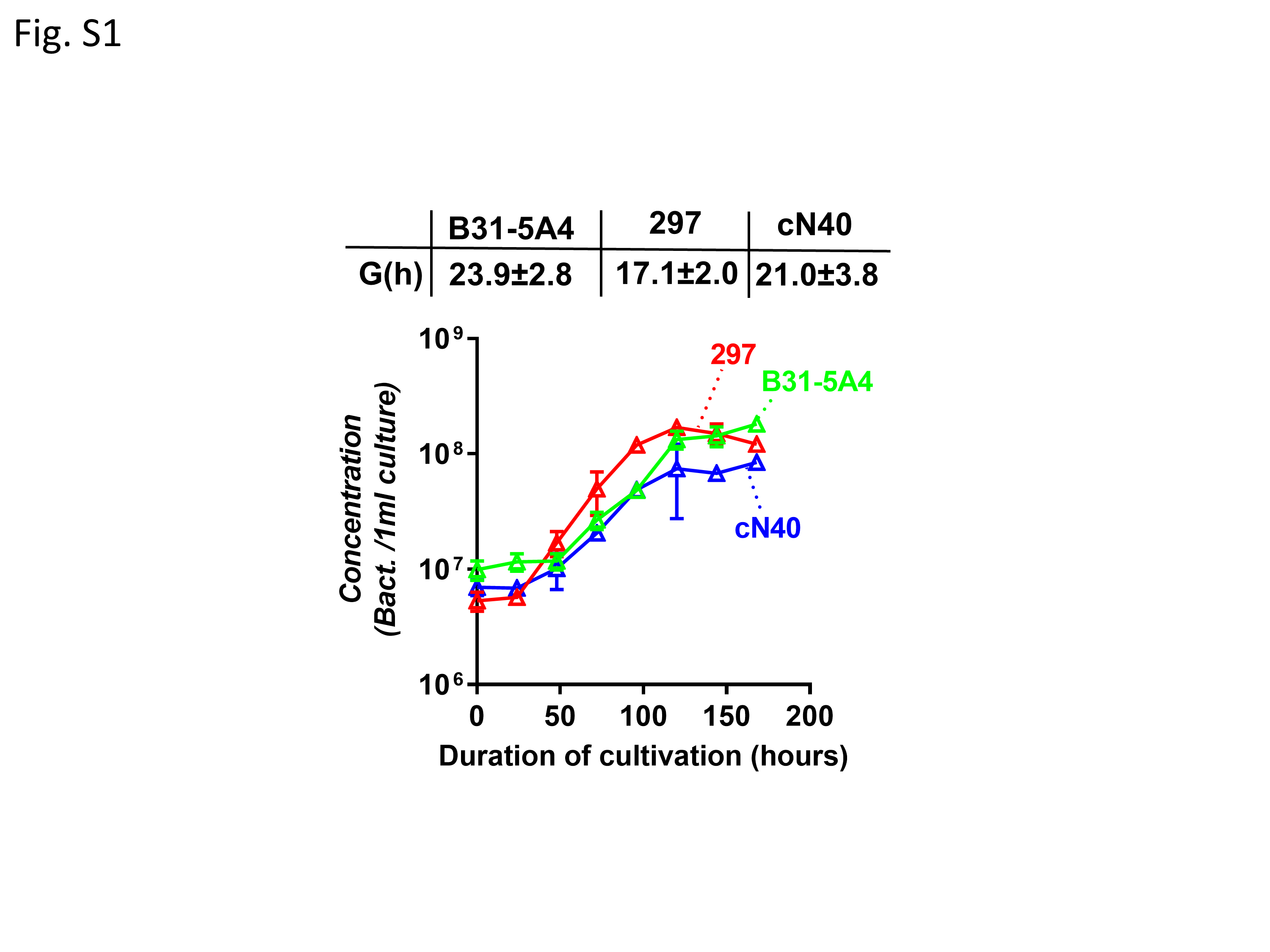

### Figure S2

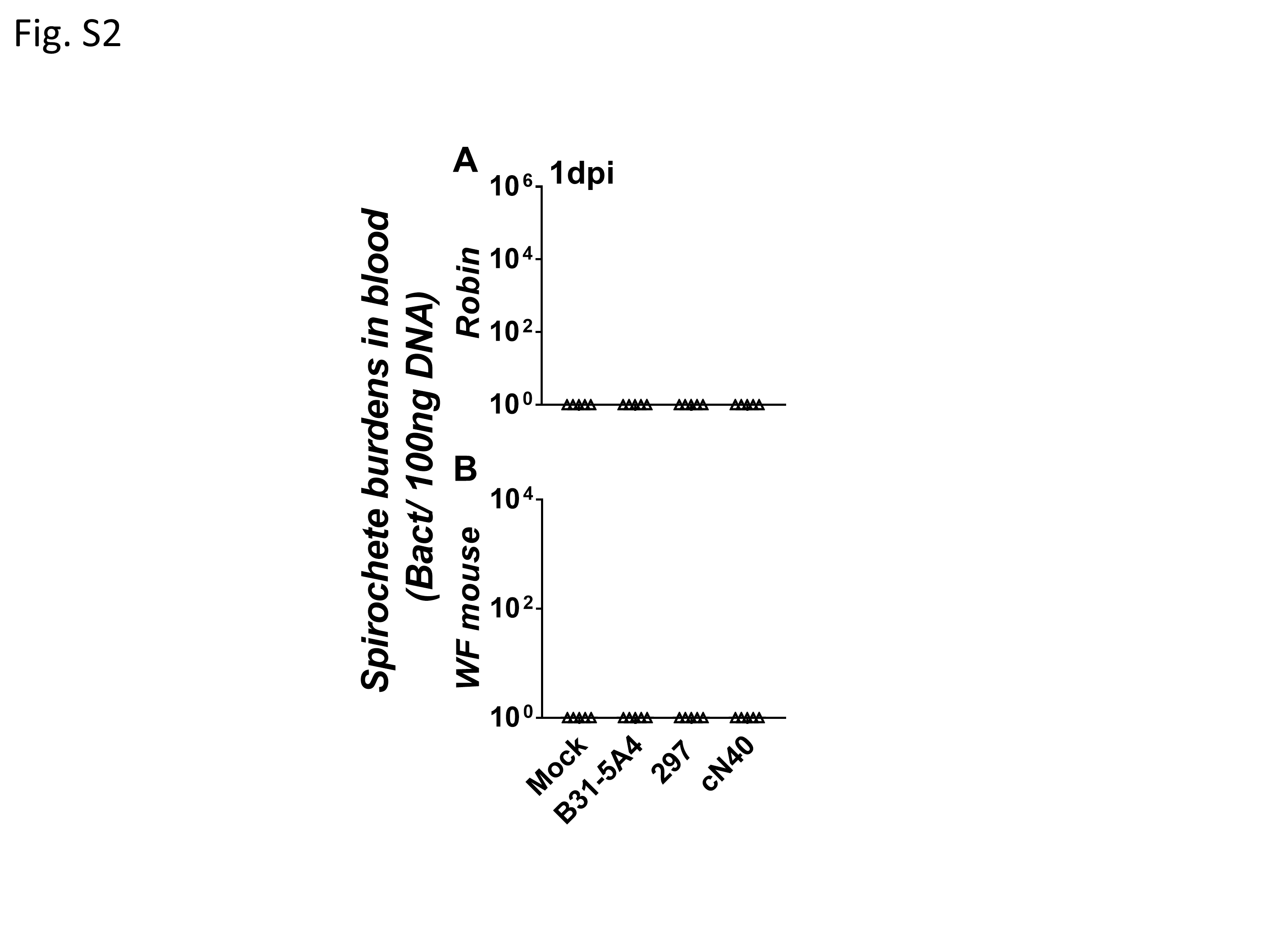

### Figure S3

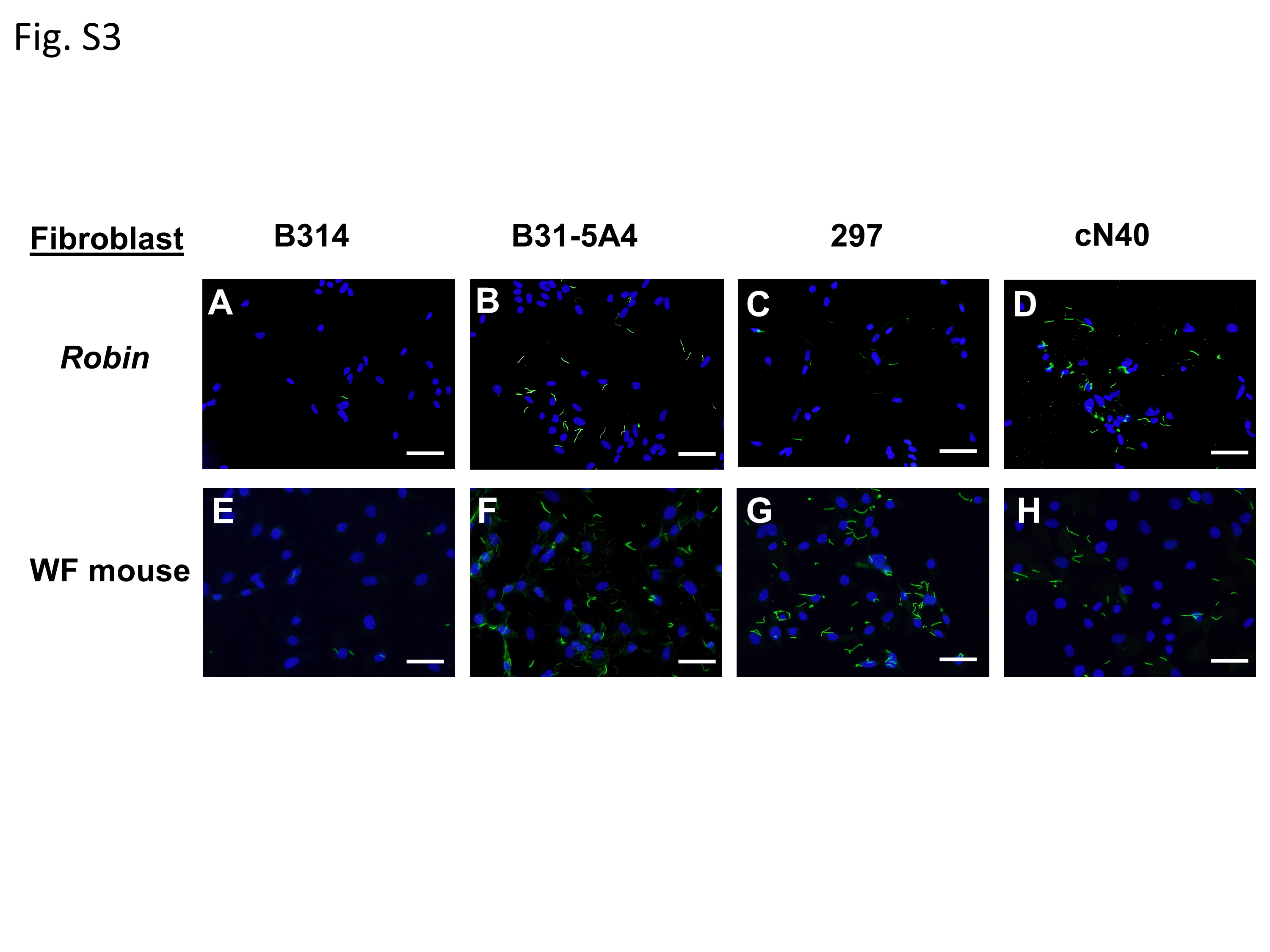

### Figure S4

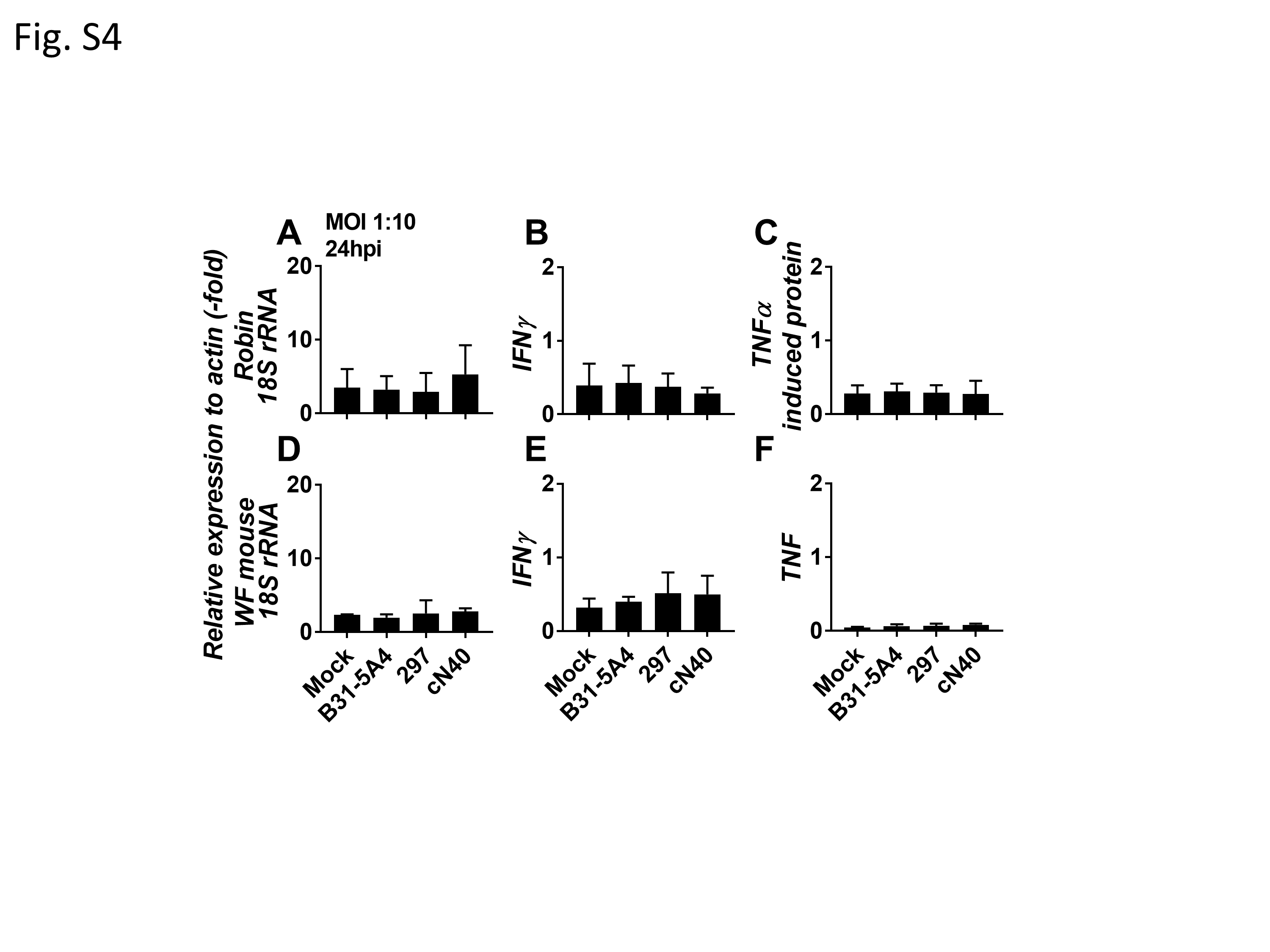

### Figure S5

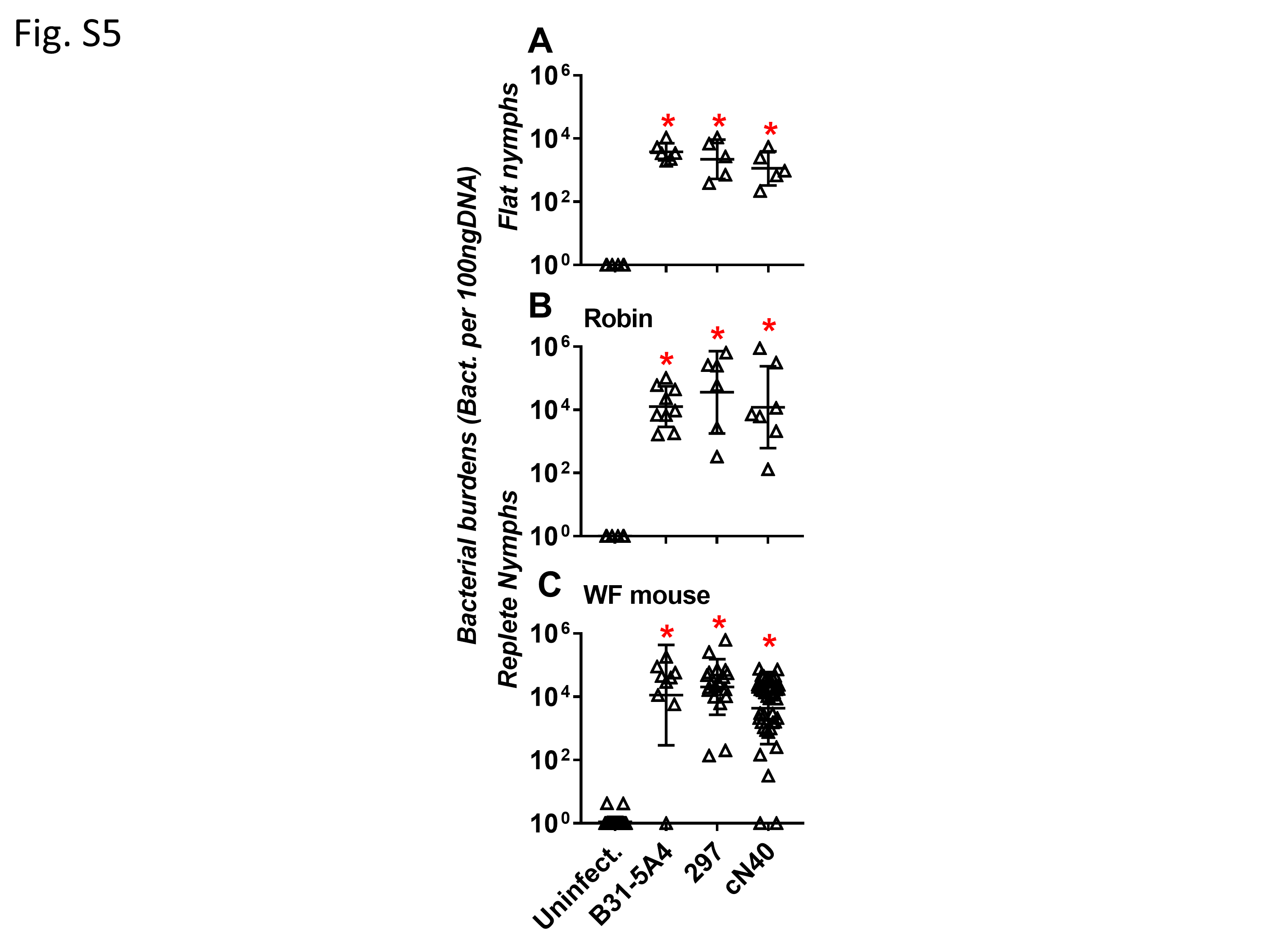
